# Hierarchical cysteine oxidation controls reversible amyloid formation in an ankyrin repeat protein

**DOI:** 10.64898/2026.08.30.748053

**Authors:** Aakriti Sethi, Hannah Darroch, Nicholas J. Magon, Grant Greene, Karina M. O’Connor, Alex D. Botha, Shelby G. Gray, Pierre de Cordovez, Rachel Coldicott, Julia Horsfield, Vanessa K. Morris, Christoph Göbl

## Abstract

The formation of amyloids, including functional amyloids, is observed for an increasing number of proteins but the molecular mechanisms that control this structural transition remain poorly understood. Here we report that the kinase inhibitor protein P18 (drP18) from *Danio rerio* (zebrafish), which contains two cysteine residues, undergoes a complex and hierarchical redox switch that strictly governs reversible amyloid formation. We identify cysteine 50 (C50) acting as a regulatory residue. Upon oxidation, C50 forms an intramolecular disulfide bond with the executioner cysteine 128 (C128), thereby blocking it. C50 can become S-glutathionylated, and upon oxidation, C128 then forms intermolecular disulfides that lead to rapid transition into amyloid fibrils. *S*-glutathionylation of C50 therefore enables amyloid formation of drP18 and the outcome is oxidant-dependent with diamide, hydrogen peroxide, peroxymonocarbonate and hypothiocyanous acid each leading to amyloid assembly with distinct kinetics and morphologies. These amyloids are fully reversible, where disulfide reduction is leading to disassembly. Whereas monomeric drP18 inhibits CDK4-mediated retinoblastoma phosphorylation, the amyloid conformation abolishes this inhibition, and reduction restores both structure and function. Expression of drP18 in zebrafish embryos yields Congo red-positive, oxidation-dependent aggregates *in vivo*. Together, our findings show that a regulatory cysteine controls an executioner cysteine to induce reversible, functional amyloid formation, revealing that proteins can encode sophisticated mechanisms to control amyloid assembly.

**Significance:** The functions of proteins are based on their three-dimensional shapes, and some can switch into alternative structures called amyloids, some of which are disease-related and others appear to have biological functions. We show that a zebrafish protein, drP18, can transition into an amyloid fold, driven by a sophisticated two-cysteine oxidation mechanism. drP18 carries two distant cysteine residues that respond differently to oxidative conditions, producing a locked form or amyloid-like aggregates depending on the conditions. Unlike disease-associated amyloids, this transition is fast, chemically controlled, and fully reversible. These results reveal how two distant cysteine residues can work together as a chemical switch to determine protein structure, suggesting that oxidation-driven amyloid formation can act as a more general mechanism of protein regulation.

## Introduction

The formation of amyloid structures involves the assembly of soluble protein building blocks into fibrillar aggregates characterized by an intermolecular cross-β sheet architecture (*1–4*). Amyloid fibrils challenge the paradigm that a protein adopts a single native structure, and they remain difficult to characterize using established structural biology approaches (*5*, *6*). Studies of disease-associated amyloids, including amyloid-β, α-synuclein and tau, have revealed complex aggregation pathways involving primary and secondary nucleation events, structural polymorphism and strong dependence on experimental conditions (*2*, *7*, *8*). A number of amyloid forming proteins have been discovered where their transition into fibrillar structures implies functional roles such as serving as interaction scaffolds (*9*) and storage systems (*10*), actively regulating biological pathways (*9*, *11–14*). The onset and the progression into amyloid structures are particularly difficult to describe (*15*). Detailed studies revealed a complex interplay between fluid dynamics and the occurrence of primary and secondary nucleation events that amplify amyloid growth (*16–19*). As a consequence, amyloid formation depends strongly on specific conditions that are frequently difficult to reproduce, even for highly purified protein preparations (*20*, *21*).

We recently discovered a novel amyloid formation mechanism where the small, globular and all-α helical human protein p16^INK4A^ (p16) forms typical β-sheet based amyloid structures strictly upon oxidation (*22*, *23*). p16 is a cell-cycle regulator and as a member of the INK4 kinase inhibitor family of tumor suppressors, it is frequently mutated in cancers (*24*). The monomeric protein adopts an ankyrin-repeat fold, consisting of helix–loop–helix motifs that stack to form an elongated, linear architecture (*25*, *26*). We have demonstrated that although p16 is stable under normal conditions, it undergoes amyloid conversion following oxidation of its single cysteine residue, which promotes the formation of intermolecular disulfide-linked dimers that subsequently assemble into amyloid fibrils (*22*). Remarkably, this process is fully reversible through reduction of the disulfide bond, revealing a direct link between cysteine redox chemistry and amyloid formation (*23*).

The oxidation-dependent transition of p16 into amyloid structures raises the possibility that redox-regulated amyloid formation is a shared property of INK4 family members; they harbor conserved ankyrin-repeat architecture but differ in cysteine content. Human p15, p16 and p18 contain one cysteine residue whereas p19 lacks this amino acid. The zebrafish (*Danio rerio*) INK4 orthologue drP18 contains two cysteines positioned within a five-repeat ankyrin scaffold. The disulfide bond chemistry of a single-cysteine protein is limited to intermolecular bonds. Here, we report that drP18 forms amyloid structures upon oxidation by an unexpectedly complex hierarchical interplay between the two functionally distinct cysteine residues. We identify C50 as a regulatory cysteine residue. Upon oxidation, C50 can form an intramolecular disulfide bond with C128, which prevents amyloid formation. C50 can also undergo *S*-glutathionylation and upon oxidation, C128 is then available to form intermolecular disulfide-linked dimers that serve as building blocks for subsequent fibril assembly. drP18 amyloid formation is therefore regulated by competition between intra- and intermolecular disulfide bond formation. Different cysteine-reactive reagents, including the model oxidant diamide and physiological species such as hydrogen peroxide, peroxymonocarbonate and hypothiocyanous acid, produce distinct amyloid formation kinetics and characteristic amyloid conformations. Importantly, depending on the type of oxidant, these concerted structural transitions are reversible through disulfide bond reduction, thereby implying a fully redox-regulated amyloid formation system that switches between two stable structural states with distinct functional impact. We further demonstrate oxidation and aggregation of drP18 in zebrafish, supporting a potential physiological relevance of this novel mechanism. Together, our functional analysis describes a hierarchical redox switch in which a regulatory cysteine controls the activity of an executioner cysteine, revealing that proteins can encode sophisticated mechanisms to control amyloid assembly.

## Results

### drP18 forms oxidant-specific amyloid-like structures

First, we confirmed the zebrafish P18 (drP18) protein sequence in our experimental fish model (WIK strain) by sequencing the *cdkn2c* locus. We identified a single point mutation in exon 2 compared to the deposited protein UniProt entry (ID: B0UXZ0), yielding an L7I substitution. We therefore used this L7I variant for further experiments, hereafter referred to as the wild-type. drP18 is an 18.7 kDa monomeric protein consisting of five ankyrin repeats containing two cysteine residues in the second and fourth repeat, and their Cα atoms are 21.2 Å apart (Fig. 1*A*). To test whether drP18 forms oxidation-induced amyloid structures similar to human p16, we performed experiments with the recombinantly expressed and purified protein.

**Fig. 1.**
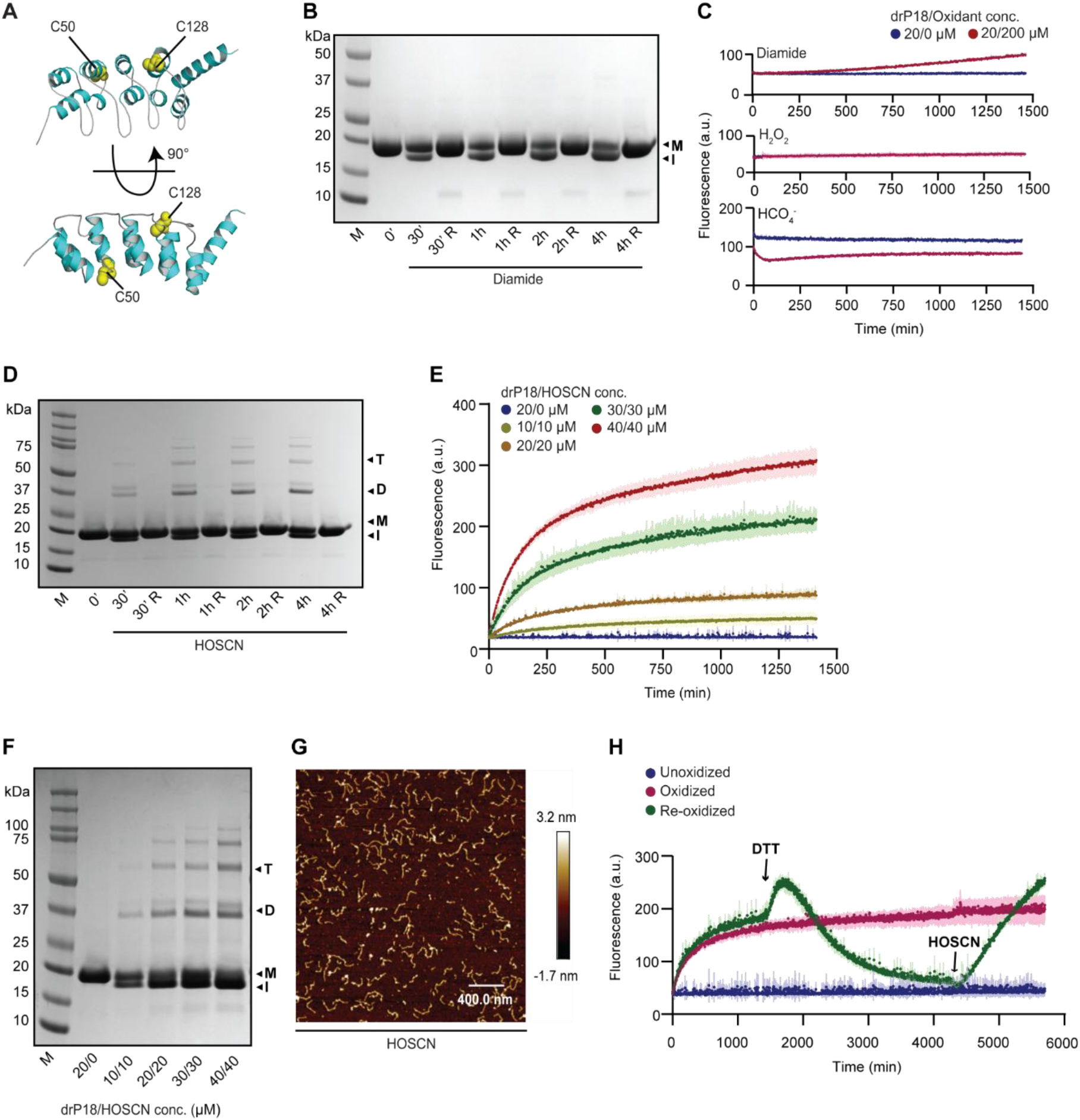
drP18 forms oxidant-specific amyloids. (A) Structural model of drP18 (AF-B0UXZ0-F1) highlighting the heavy atoms of the two cysteine residues C50 and C128 as yellow spheres (Cα– Cα distance = 21.2 Å). (B) SDS-PAGE analysis of 20 µM drP18 following oxidation with 200 µM diamide over a time course of 4 h at 25 °C. The label M indicates the monomeric band, I indicate formation of an intramolecular disulfide bond. Addition of the reducing agent DTT (R) reduces the disulfide bond. (C) Thioflavin-T fluorescence (ThT) kinetic assays of drP18 and oxidation by diamide (top), H_2_O_2_ (center) and HCO_4_^-^ (bottom). 20 µM protein was oxidized with 200 µM oxidant (1:10 protein to oxidant ratio). (D) SDS-PAGE analysis of 20 µM drP18 (M) shows formation of I, intramolecular disulfide bond and D, dimer following oxidation with 20 µM HOSCN over a time course of 4 h at 25 °C. Addition of the reducing agent DTT breaks the disulfide bonds. (E) ThT kinetic assays of drP18 (10–40 µM) oxidized with HOSCN at a fixed 1:1 molar ratio at 25 °C. (F) SDS-PAGE analysis of drP18 across a concentration series at a 1:1 protein-to-HOSCN ratio over a 4 h time course at 25 °C. M, I, refer to species described in (B), D and T refer to dimeric and trimeric species, respectively. (G) AFM analysis of 20 µM drP18 oxidized with 20 µM HOSCN for 24 h at 25 °C. Scale bar: 400.0 nm. (H) Thioflavin-T fluorescence kinetics assay of drP18 (20 µM) oxidized with HOSCN (20 µM). Addition of 200 µM DTT after 1430 minutes (indicated by left arrow) causes reduction of fluorescence to baseline levels and subsequent re-addition of 420 µM HOSCN at 4400 minutes (indicated by right arrow, excess to quench DTT) leads to an increase of the ThT fluorescence.

We first tested a panel of oxidants including diamide, a cysteine-specific oxidant (*27*), and the physiological oxidants hydrogen peroxide (H_2_O_2_) and peroxymonocarbonate (HCO_4_ ^-^, generated from H_2_O_2_ and bicarbonate). Initial protein oxidation experiments were performed at previously established protein-to-oxidant ratios of 1:10 (*23*). Diamide produced an intra-molecular disulfide bond upon oxidation for 30 min to 4 hours, as evidenced by faster migrating bands in SDS-PAGE analysis. These were fully reversed by the addition of the disulfide reductant dithiothreitol (DTT) (Fig. 1*B*). No dimers were detected during this experiment, and the results were similar for H_2_O_2_ and HCO_4_ ^-^ (Fig. S1 *A* and *B*). None of these oxidants led to an increase in fluorescence in assays using Thioflavin T (ThT, Fig. 1*C*), a dye widely used to detect amyloid structures (*22*, *28*). This suggested that intra-molecular disulfide formation does not drive transition into fibrillar structures.

A different outcome was observed when testing the immune-derived oxidant hypothiocyanous acid (HOSCN). Different protein-to-oxidant ratios were tested, and an increase in ThT fluorescence was observed for a 1:1 equimolar mixture of protein and oxidant (Fig. S1*C*), but not for other ratios. We previously confirmed that the oxidants tested do not interfere with the assay (*29*). SDS-PAGE analysis displayed the formation of dimers and higher-order species upon incubation with equimolar HOSCN levels for 30 min to 4 hours (Fig. 1*D*), and this was fully reducible upon addition of DTT. ThT assays indicated rapid amyloid formation at protein concentrations in the 10–40 μM range in the presence of equimolar oxidant (Fig. 1*E*) and similar higher order inter-molecular species were observed in SDS-PAGE (Fig. 1*F*). Atomic force microscopy (AFM, Fig. 1*G*) and electron microscopy (EM, Fig. S1*G*) provided complementary morphological evidence for amyloid fibril formation. When changing protein-to-oxidant ratios for the other oxidants, including a 1:1 stoichiometry, no significant differences were observed between the different conditions as monitored by ThT assays (Fig. S1 *D* and *E*) and SDS-PAGE analysis (Fig. S1*F*). AFM analysis showed no presence of aggregates or amyloid-like structures (Fig. S1*H*).

Previously, amyloid formation of human p16 was found to be reversible upon addition of reducing agents that break the inter-molecular disulfide bond (*23*). We therefore assessed whether drP18 amyloid structures are reversible. Upon addition of DTT during ThT assays, the drP18 amyloids were disassembled, and they again formed upon re-addition of HOSCN (1:1 protein-to-HOSCN ratio, Fig. 1*H*, Fig. S2*A*). Consistent with the ThT data, reduction was accompanied by a loss of dimeric and higher-order species on SDS-PAGE (Fig. S2*B*). When analyzing the HOSCN-mediated oxidation product by mass spectrometry, besides the presence of the intramolecular C50-C128 disulfide bond, the predominant intermolecular disulfide linkage involved C128 residues (C128–C128), whereas a C50–C50 disulfide linkage was not observed under these conditions (Fig. S2*C*). The m/z values for the parent ions and the daughter fragment ions that were used to quantify each peptide in LC-MS/MS experiment are listed in Table S1.

In parallel, we confirmed proper folding of our protein construct by ^1^H solution NMR spectroscopy, where uniformly labelled ^15^N drP18 displayed a well-dispersed, structured ^1^H^15^N HSQC spectrum (Fig. S3*A*, top). When the sample was oxidized with diamide for 24 h, an intensity reduction, mostly of the dispersed peaks, was observed with additional peaks appearing in the unstructured region (Fig. S3*A*, center). Likely, these changes indicate the transition of a fraction of the protein into a species with an intra-molecular disulfide bond that is partially unstructured that lacks aggregation tendency. Addition of DTT reversed the spectrum to the initial, fully reduced state, similar to what was observed by SDS-PAGE (Fig. S3*A*, bottom). Oxidation with equimolar concentrations of HOSCN led to a similar intensity reduction of structured peaks, likely with higher-order species formed that are too large to yield signals (Fig. S3*B*, top, center). Reduction of the disulfide bond with DTT again led to recovery of the reduced spectrum (Fig. S3*B*, bottom).

These results demonstrate that most oxidation conditions tested for drP18 lead to intra-molecular disulfide-bonded species, which implies large structural flexibility of the protein. The oxidation with a specific 1:1 protein-to-oxidant ratio with HOSCN is able to generate cross-linked multimers that further transition into fibrillar structures with typical amyloid-like features. HOSCN is a cysteine-specific, fast oxidant and we hypothesize that the oxidation rate impacts the outcome of cross-linking events.

### C50 acts as a regulatory residue controlling drP18 fibril formation

We next examined the reactivity of the two cysteine residues of drP18 by measuring their redox potentials. For this, we used solution NMR spectroscopy with which we previously obtained the backbone chemical shift assignments to identify individual amino acids (*30*). The secondary chemical shifts highlight the presence of helix-loop-helix motifs, consistent with the structural model of drP18. The redox potential at pH 7.4 was determined using uniformly ^15^N-labeled protein and a reduced and oxidized glutathione buffer system, as glutathione is the predominant intracellular thiol and thus a physiologically relevant redox couple. A ^1^H^15^N 2D-HSQC solution NMR spectrum was recorded in the presence of 0.4 mM reduced glutathione (GSH) to measure the reduced reference state. A redox titration was then performed, in which the GSH concentration was kept constant and the oxidized glutathione (GSSG) concentration was gradually increased. After each titration step, a ^1^H^15^N 2D-HSQC spectrum was measured and residue-specific chemical shift changes were monitored (Fig. 2*A*). The available assignments for the reduced form of drP18 were used to follow the decaying peak intensities during the titration series, allowing for calculation of the redox potential via the Nernst equation (31). Intensities of the regions with well-dispersed peaks were fitted to a decaying sigmoidal curve, from which the midpoint potential (E°’) was obtained. Affected peaks were fitted individually to a Boltzmann decay function using the known reference values for glutathione (*31*). A subset of peaks including C50 exhibited the earliest changes upon titration, yielding the average redox potential of -153.7 ± 3.3 mV (orange, Fig. 2*B*). Another group of residues transitioned into the oxidized state at -134.3 ± 0.7 mV (magenta, Fig. 2*B*). A subset of peaks, including C128, was affected by both cysteine oxidation events (Fig. S4*A*). These results demonstrated differential redox reactivity of the two cysteine residues towards glutathione, with C50 being more readily oxidized than C128.

**Fig. 2.**
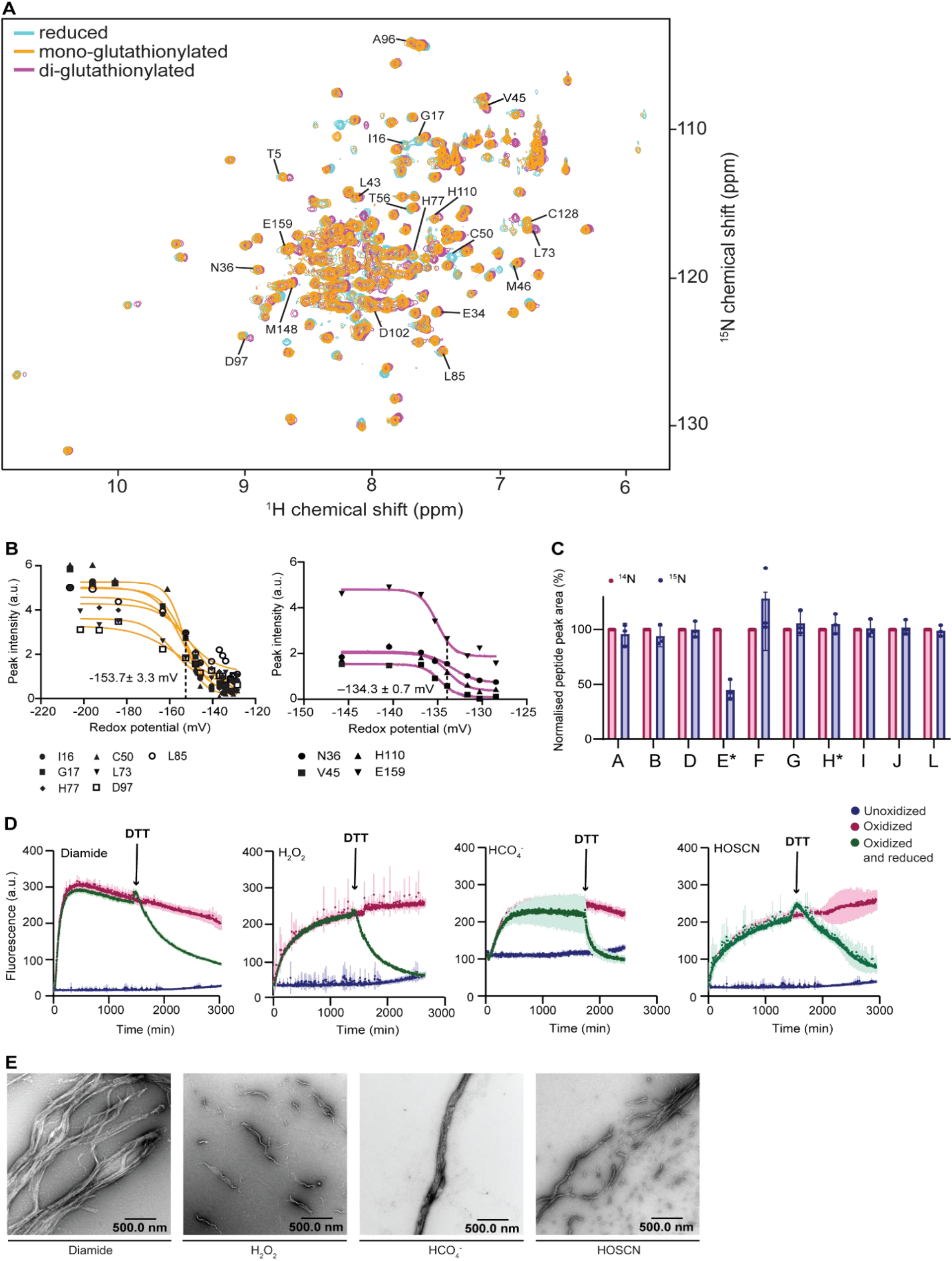
C50 is the regulatory cysteine controlling drP18 fibril formation. (A) ^1^H^15^N HSQC solution NMR spectrum of drP18 in the reduced state (cyan) and at an intermediate stage of S-glutathionylation (orange), and following complete S-glutathionylation (magenta). Residues undergoing significant chemical-shift perturbations are labelled. (B) Sigmoidal fitting curves of peak intensities of residues impacted by the first observed glutathionylation of C50 are shown in orange, residues impacted by the later observed glutathionylation of C128 are shown in magenta. The individual amino acid assignments are displayed in the legend underneath the graph. (C) Glutathione-treated ^15^N-labelled drP18 was mixed with reduced ^14^N-labelled drP18 as an internal reference standard, alkylated with NEM, digested with trypsin, and analyzed by LC-MS. Peptides are described in Fig. S4C and peak areas were quantified and normalized to the corresponding ^14^N peptide. Cysteine-containing peptides are indicated by an asterisk (*). Reduced abundance of an NEM-labelled ^15^N peptide relative to the corresponding ^14^N reference peptide indicates S-glutathionylation of that cysteine residue. (D) ThT kinetic assays of drP18 C50–SSG (20 µM) treated (left to right) with (magenta) or without (blue) 200 µM diamide, 200 µM H_2_O_2_, 200 µM HCO_4_^-^ and 20 µM HOSCN. Addition of 2 mM DTT at 24 h (indicated by arrow, green) causes reduction of fluorescence close to baseline levels. Four replicates were measured per sample and the average is plotted as a solid line, and error bars represent the standard deviation. (E) Representative electron microscopy (EM) images of oxidant-treated drP18 C50–SSG. The oxidants diamide, H_2_O_2_, HCO_4_^-^, and HOSCN induce morphologically distinct amyloid-like assemblies. Scale bar: 500 nm.

The differences in reactivity and reaction kinetics are large enough for glutathione conditions to generate a species in which C50 is predominantly oxidized, while C128 remains reduced. Using the wild-type protein, we produced the single *S*-glutathionylated protein species (drP18 C50– SSG, as confirmed by NMR spectroscopy, Fig. S4*B*) to test its behavior upon oxidation. We further confirmed the site specificity of *S*-glutathionylation by mass spectrometry following tryptic digestion of drP18, enabling identification and quantification of modifications at the peptide level. To distinguish modified from unmodified peptides, we generated two drP18 WT samples, reduced protein at natural nitrogen abundance (^14^N) and uniformly ^15^N-labelled protein that had been *S*-glutathionylated as described above. Mixing of the two isotopically distinct samples for combined analysis allowed direct comparison of corresponding peptides, which co-eluted during LC-MS analysis but were readily distinguished by their mass difference. Prior to digestion, both the reduced ^14^N and glutathionylated ^15^N-labelled proteins were treated separately with 10 mM *N*-ethylmaleimide (NEM) to alkylate all remaining free cysteine residues. Because cysteines occupied by glutathione are inaccessible to NEM, *S*-glutathionylation results in a corresponding decrease in the abundance of the NEM-labelled peptide. Following NEM treatment, proteins were digested with trypsin, followed by LC-MS analysis where peptide peak areas were extracted and quantified. Comparison of the ^15^N-labelled glutathionylated sample with the reduced ^14^N control revealed a pronounced loss of the NEM-labelled peptide containing C50, whereas little change was observed for the peptide containing C128. These data demonstrate that *S*-glutathionylation occurs predominantly at C50 (Fig. 2*C*, Fig. S4*C*). The peptide sequence and theoretical m/z values used for quantification of the glutathionylated peptide are provided in Table S2.

Selective glutathionylation of C50 leaves C128 as a free cysteine residue and we exposed drP18 C50–SSG to different oxidants to determine the impact of oxidation. Incubation with diamide, H_2_O_2_, HCO_4_^-^ and HOSCN all readily led to formation of amyloids as probed by ThT assays (t_1/2 (Diamide)_ = 75.3 ± 7.7 min, t_1/2 (H2O2)_ = 282.7 ± 9.4 min, t_1/2 (HCO4-)_ = 158.1 ± 9.5 min, t_1/2 (HOSCN)_ = 207.5 ± 20.3 min). All samples show loss of ThT intensity upon DTT addition, suggesting disassembly of the amyloid species and therefore reversible amyloid transitions (Fig. 2*D*, Fig. S4*D*). For each oxidant, the ThT fluorescence increased rapidly and reached a plateau without a detectable lag phase, suggesting accelerated nucleation and a shortened amyloid growth phase compared to the previous experiments. In addition, electron microscopy revealed large fibrillar structures in all oxidant-treated drP18 C50–SSG samples (Fig. 2*E*) with some individual characteristics. Diamide treatment resulted in long, bundled fibrillar assemblies, while H_2_O_2_ produced shorter, more fragmented fibrils with less extensive bundling. HCO_4_^-^ generated thick fibrillar structures, and HOSCN promoted curved and heterogeneous filaments. Together, these observations reveal that C50 acts as a key switch allowing various oxidants to trigger C128-dependent intermolecular crosslinking and amyloid formation. Blocking of C50 therefore strikingly changes the oxidation response of drP18 and leads to a major structural rearrangement.

To determine whether C50 undergoes reversible glutathionylation, we analyzed drP18 by intact-protein mass spectrometry. Glutathionylated ¹⁵N-labelled drP18 produced a predominant species corresponding to the addition of one glutathione moiety, with a measured mass of 19,247 Da, in close agreement with the predicted mass of 19,245.5 Da. Treatment with DTT restored the protein to its unmodified mass of 18,941 Da (predicted, 18,940.4 Da), demonstrating that the glutathione modification is reversible (Fig. S4*E*).

To further investigate whether glutathione interaction contributes to cysteine-dependent modification, we examined the interaction of GSH with the cysteine-free double mutant (drP18 C50S C128S). Addition of a 10-fold molar excess of GSH resulted in subtle chemical shift perturbations in a subset of amino acid resonances (Fig. S5), indicating weak non-covalent interaction of glutathione with drP18 even in the absence of cysteine residues. These observations suggest that GSH can transiently associate with the protein scaffold, which may facilitate C50 glutathionylation in WT drP18.

### The C128 executioner cysteine drives amyloid formation

Having described a regulated two-cysteine mechanism of the wild-type protein leading to amyloid formation, we next further confirmed the roles of the cysteines by studying the recombinantly expressed cysteine variants drP18 C50S, drP18 C128S, and the double cysteine mutation construct drP18 C50S C128S. The proteins were purified and exposed to several oxidants.

The drP18 C50S protein formed a dimer (∼37.4 kDa) upon oxidation with diamide, and this dimer was largely non-reducible, showing only partial reduction upon DTT treatment in SDS-PAGE (Fig. 3*A*). The oxidation led to a transition into amyloid according to ThT assays, with increasing kinetic rates at higher oxidant concentrations (half-time of amyloid formation as fast as t_1/2_ = 36.3 ± 0.3 min at a protein-to-oxidant ratio of 1:20, Fig. 3*B*). In contrast, H_2_O_2_ treatment did not produce detectable dimer formation within the 4 h incubation period and ThT fluorescence increased only gradually over time, indicating relatively slow amyloid assembly under these conditions (Fig. 3 *C* and *D*). HCO_4_^-^ promoted the slow formation of dimeric species and a concentration-dependent increase in ThT fluorescence with faster formation rates at higher oxidant concentrations (Fig. 3 *C* and *D*). HOSCN addition induced dimer formation and led to ThT fluorescence exclusively under equimolar conditions (protein-to-oxidant ratio of 1:1), similar to what was observed for the wild-type protein. Higher dr P18 C50S protein and oxidant concentrations at a fixed protein-to-oxidant ratio (1:10, except 1:1 for HOSCN) resulted in progressively greater accumulation of dimeric species (Fig. S6*A*) as well as increased ThT fluorescence (Fig. S6*B*). Addition of DTT to drP18 C50S amyloids resulted in only partial loss of ThT fluorescence in diamide- and HOSCN-treated samples, suggesting that the resulting higher-order structures were not fully reversible. In contrast, HCO_4_^-^ -induced assemblies were fully reversed by DTT, with ThT fluorescence returning to baseline levels (Fig. S6*C*). Electron microscopy revealed that the type of oxidant strongly influences the morphology of amyloid assemblies formed by oxidation of drP18 C50S. Diamide treatment produced compact, clustered aggregate structures with relatively dense and amorphous morphology. In contrast, H_2_O_2_ induced highly ordered, elongated fibrillar bundles. HCO_4_ ^-^ generated long, thick fibrillar structures with occasional branching, whereas HOSCN produced more curved and heterogeneous filaments with less bundling (Fig. 3*E*).

**Fig. 3.**
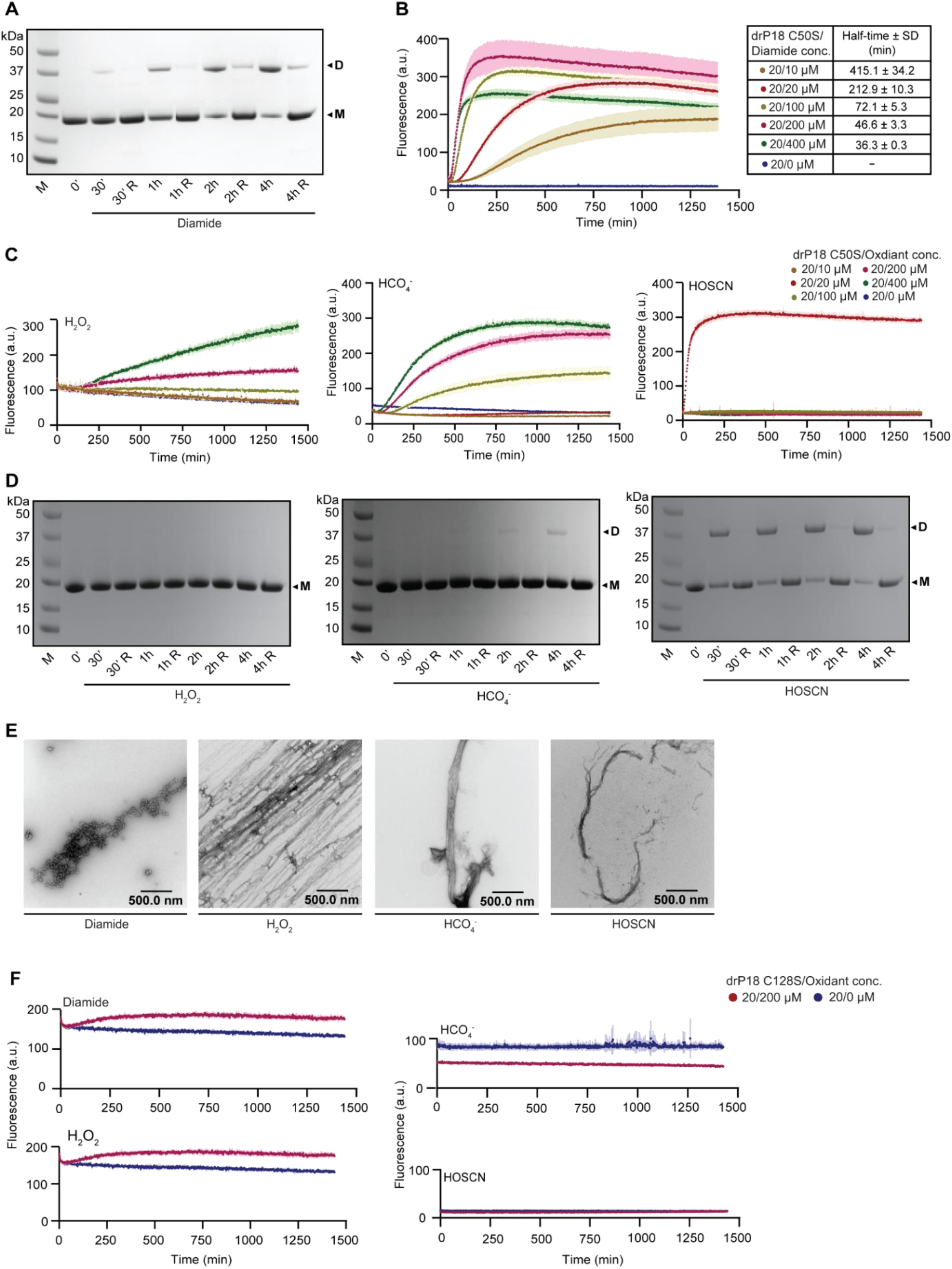
Oxidation of drP18 C128 leads to amyloid formation. (A) SDS-PAGE analysis of 20 µM drP18 C50S (M) shows formation of D, dimer following oxidation with 200 µM diamide over a time course of 4 h. Addition of the reducing agent DTT resulted in partial reduction of disulfide bond intensity. (B) Thioflavin-T fluorescence kinetics assay of drP18 C50S (20 µM) oxidized with different concentrations of diamide (0-400 µM). (C) Thioflavin-T fluorescence kinetics assay of drP18 C50S (20 µM) oxidized with different oxidants at concentrations ranging from 10–400 µM. (D) SDS-PAGE analysis of 20 µM drP18 C50S (M) following oxidation with 200 µM H_2_O_2_ (left), 200 µM HCO_4_^-^ (center) and 20 µM HOSCN (right) over a time course of 4 h, where a D, dimer is observed, which is partially reduced upon addition of DTT. (E) Representative electron microscopy (EM) images of drP18 C50S assemblies formed under different oxidizing conditions. Distinct amyloid morphologies are observed following treatment with diamide (left), H_2_O_2_ (center left), HCO_4_^-^ (center right), and HOSCN (right). Scale bar: 500 nm. (F) Thioflavin-T fluorescence kinetics of drP18 C128S (20 µM) following oxidation with diamide (200 µM), H_2_O_2_ (200 µM), HOSCN (20 µM), or HCO_4_^-^ (200 µM) at 25 °C. No increase in ThT fluorescence was observed over the 24 h time course under any condition.

Experiments with drP18 C128S revealed faint dimer band formation following oxidation for up to 4 h with diamide, H_2_O_2_, and HOSCN, whereas HCO_4_ ^-^ induced a more pronounced dimeric species (Fig. S7*A*). Despite dimer formation, no amyloid-type assemblies were detected, as indicated by the absence or only marginal increase of ThT fluorescence (Fig. 3*F*, Fig. S7 *B* and *C*) and the lack of fibrillar structures visualized by AFM analysis (Fig. S7*D*).

The double cysteine mutant did not exhibit dimer formation or an increase in ThT fluorescence with any of the oxidants tested, supporting the cysteine oxidation driven amyloid formation mechanism (Fig. S8 *A* and *B*). However, AFM imaging of the double cysteine mutant revealed fibril-like and entangled morphologies in the unoxidized state, suggesting that removal of both cysteines compromises the stability of the protein and promotes spontaneous, non-oxidation driven aggregation (Fig. S8*C*). We characterised the stability of the protein cysteine variants using differential scanning fluorimetry (DSF). The drP18 WT protein exhibited a relatively low melting temperature (Tₘ = 38.7 ± 0.2 °C). Mutation of either cysteine residue further reduced thermal stability, with Tₘ values of 37.3 ± 0.1 °C for drP18 C50S and 34.2 ± 0.2 °C for drP18 C128S, while the double mutant showed the lowest stability (Tₘ = 32.8 ± 0.3 °C). We further tested some cysteine to alanine mutations which had a similar reduction in melting temperature (C50A Tₘ = 34.7 ± 0.4 °C and C50A C128A Tₘ = 32.6 ± 0.3 °C, data not shown). This progressive decrease highlights the crucial role of both cysteines in maintaining structural integrity of the monomeric, folded conformation (Fig. S8*D*).

These experiments establish that the transition of drP18 into amyloid structures is governed by the C128 residue. Interestingly, the disulfide-reduction driven dissociation is less prevalent for drP18 C50S amyloids induced by diamide and HOSCN. This could be due to intrinsic structural differences such as disulfide accessibility and amyloid stability, or the impact of formation kinetics leading to different dissociation properties.

### Amyloid formation prevents inhibition of CDK4 by drP18

The transition of drP18 into amyloid assemblies might impact its role as a kinase inhibitor, and we next examined whether this structural conversion alters its ability to inhibit CDK4. To address this, we performed an *in vitro* kinase assay using human recombinant GST-CDK4/cyclin D1 and an Rb substrate, with phosphorylation of Rb at Ser807/811 as a measure of CDK4 activity (*23*). Robust phosphorylation was observed in the positive control lacking inhibitors, whereas no signal was detected in the absence of CDK4, confirming the specificity of the assay (Fig. 4*A*). In its native, monomeric state, drP18 efficiently inhibited CDK4 activity, consistent with its structural similarity to human INK4 proteins (∼65% sequence similarity to human p18). In contrast, amyloid assemblies generated by oxidation with HOSCN (protein-to-oxidant ratio of 1:1) allowed Rb phosphorylation, indicating a loss of inhibitory function upon aggregation. Notably, reduction of the oxidized fraction with 10 mM DTT for 16 h partially restored CDK4 inhibition (Fig. 4*A*). We next performed kinase assays using C50-glutathionylated drP18. In its monomeric form, drP18 C50–SSG potently inhibited CDK4 activity, as indicated by a marked reduction in phospho-Rb levels (Fig. 4*B*). However, transition into amyloids through oxidation of drP18 C50–SSG with both, diamide or peroxymonocarbonate abolished this inhibitory effect, demonstrating that modification at C50 promotes a loss of function upon oxidation. Subsequent reduction restored inhibitory activity, consistent with a fully reversible redox-dependent structural change. We further examined the drP18 C50S mutant under identical conditions (Fig. 4*C*). The protein retained CDK4 inhibitory activity comparable to that of drP18 C50–SSG. Oxidation of C50S with either diamide or HCO_4_ ^-^ oxidation similarly abolished inhibitory activity, as indicated by persistent Rb phosphorylation. Following reduction, HCO_4_ ^-^ treated drP18 C50S samples regained inhibitory function, evidenced by loss of the pRb signal, whereas diamide-treated samples remained inactive with persistent pRb detection. These findings indicate that diamide-induced drP18 C50S assemblies are not fully reversible, in contrast to the reversibility observed following HCO_4_ ^-^ mediated oxidation. This is in line with the observed amyloid disassembly through reduction in the ThT fluorescence assays (Fig. S6*C*).

**Fig. 4.**
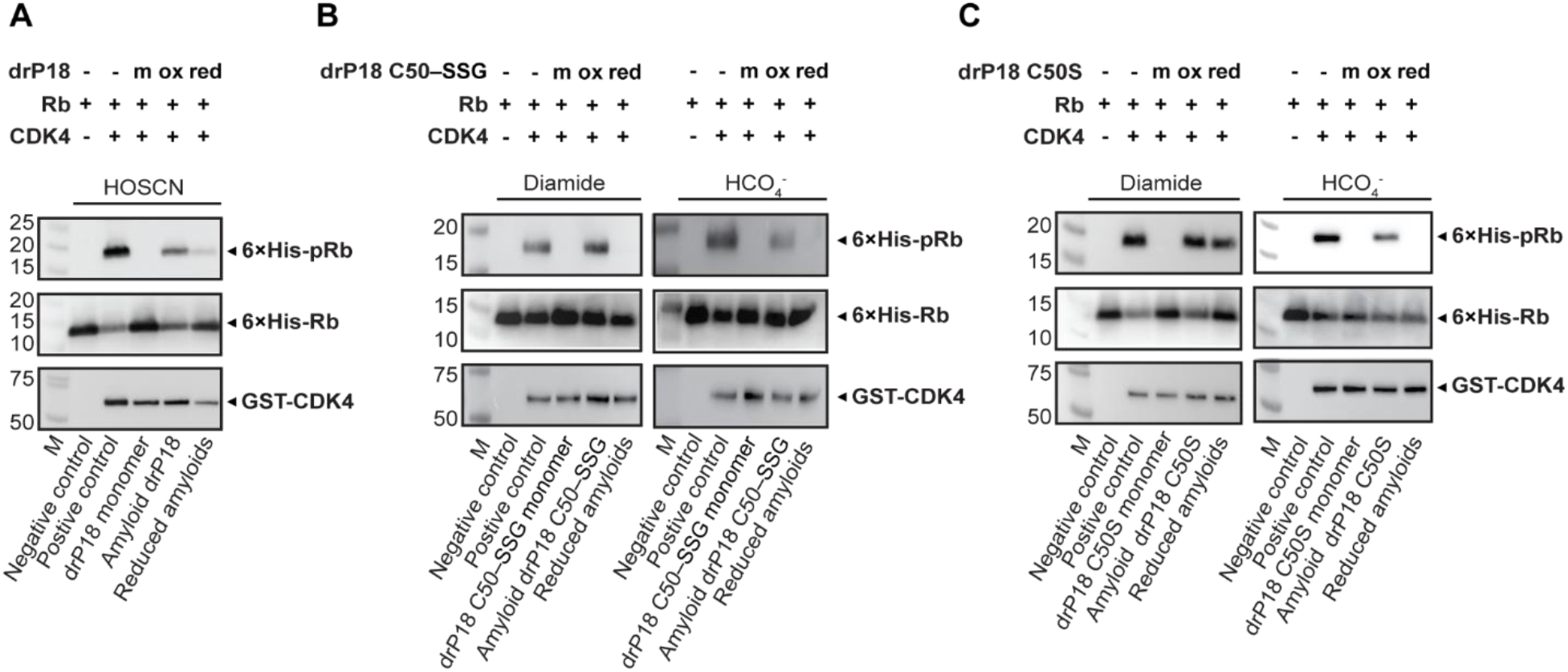
C50 glutathionylation reversibly regulates drP18 CDK4 inhibition. (A) In vitro kinase assays using recombinant GST–CDK4/Cyclin D1 and 6×His–Rb substrate in the presence of drP18 were performed. Monomeric drP18 inhibits CDK4-mediated phosphorylation of Rb, whereas amyloid assemblies generated by oxidation (HOSCN, protein-to-oxidant ratio of 1:1) allow kinase activity. Reduction of oxidized drP18 with 10 mM DTT for 16 h partially restores inhibitory function, as indicated by decreased phospho-Rb signal. (B) The kinase assays were performed with drP18 C50–SSG. The monomeric protein inhibits CDK4 activity, whereas oxidation with either diamide (left) or HCO_4_^-^ (right) abolishes this inhibitory effect and reduction restores kinase inhibition. (C) The kinase assay was performed with the drP18 C50S mutant and the monomeric protein inhibits CDK4 activity. Oxidized drP18 C50S failed to inhibit Rb phosphorylation following treatment with both diamide and HCO_4_^-^ oxidants. However, reduction restored inhibitory function only in the HCO_4_^-^ treated samples, as indicated by the loss of the pRb band, whereas the diamide-treated samples retained pRb signal after reduction, indicating that diamide-induced amyloid formation was not fully reversible. Phosphorylation of Rb (Ser 807/811) was detected by immunoblotting. Total Rb and GST–CDK4 are shown as loading controls. Negative control lacks CDK4; positive control contains CDK4 and Rb only. Representative images from two independent experiments are shown.

Together, these qualitative findings demonstrate that glutathionylation at C50 can regulate drP18 activity by promoting a reversible transition into an inactive amyloid state. Substitution of C50 disrupts this redox-sensitive switch, resulting in impaired disassembly upon reduction and highlighting the role for this residue in mediating redox-dependent functional regulation.

### Amyloid formation of drP18 eGFP variants in zebrafish embryos

We next investigated whether oxidation and amyloid formation of drP18 variants occur *in vivo* by ectopically expressing the proteins in zebrafish embryos (Fig. 5*A*). Wild-type zebrafish (WIK) were bred, and embryos were collected at the one-cell stage. Embryos were then microinjected with mRNA encoding drP18 eGFP, drP18 C50A eGFP (the C50A variant was chosen to avoid potential phosphorylation), scrambled mRNA, or eGFP alone as a control. To confirm that the eGFP fusion did not alter drP18 oxidation behavior, the drP18 eGFP constructs were also recombinantly expressed in bacteria and purified and they showed comparable properties upon oxidation to the corresponding untagged proteins (Fig. S9, *A–D*).

**Fig. 5.**
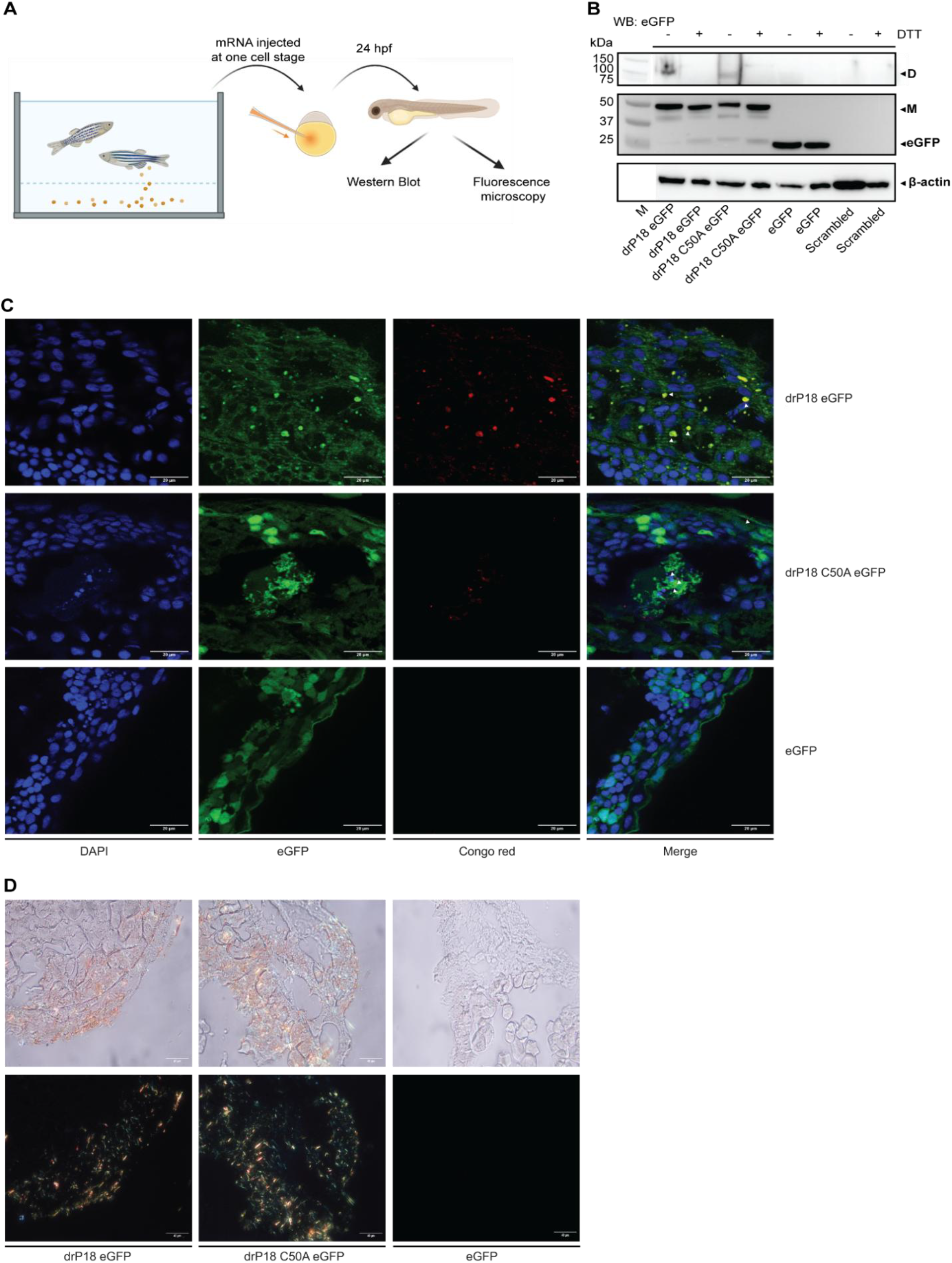
Amyloid formation of drP18 eGFP variants in zebrafish embryos. (A) Overview of the experimental workflow. Wild-type (WIK) zebrafish embryos were collected at the one-cell stage and microinjected with mRNA encoding drP18 eGFP, drP18 C50A eGFP, scrambled mRNA, or eGFP alone. Embryos were analyzed at 24 hours post-fertilization (hpf). The figure was created with BioRender. (B) Western blot of 24 hpf embryo lysates probed with anti-GFP antibody. Monomeric bands corresponding to drP18 eGFP, drP18 C50A eGFP, and eGFP are detected. A higher molecular weight band consistent with dimer formation is observed in drP18 eGFP and drP18 C50A eGFP embryos, indicating oligomerization in vivo. β-actin (∼42 kDa) was used as a loading control. (C) Representative fluorescence image of 10 µm cryosections from embryos expressing drP18 eGFP, drP18 C50A eGFP and eGFP alone respectively stained with Congo Red. Co-localization of Congo Red signal with drP18 eGFP and drP18 C50A eGFP fluorescence is observed, consistent with protein association with Congo Red-positive deposits. Scale bar: 20 µm. (D) Polarized light microscopy of Congo Red-stained embryos showing apple-green birefringence in regions corresponding to Congo Red-positive deposits, consistent with amyloid structures. Data represents n = 8 embryos per condition; representative images shown. Scale bar: 40 µm

At 24 hours post-fertilization (hpf), embryos were screened by fluorescence microscopy for eGFP expression, homogenised, and lysates were analyzed by western blot using an anti-GFP antibody. Expression of monomeric drP18 eGFP, drP18 C50A eGFP, and eGFP alone was detected, confirming efficient *in vivo* translation of the injected constructs. A lack of bands in the scrambled control suggests specificity of the used antibody. Faint but visible higher molecular weight species consistent with the molecular weight of dimers were observed in embryos expressing drP18 eGFP and drP18 C50A eGFP; both disappeared under reducing SDS-PAGE conditions (Fig. 5*B*). These findings indicate that the wild-type and the C50A drP18 variant undergo *in vivo* dimerization under physiological embryonic conditions that is reversible by treatment with reducing agents, suggesting a cysteine-dependent linkage. The oxidation is likely driven by generation of endogenous oxidants, which have been reported previously to be present at high levels during embryonic development (*32*).

To assess potential amyloid formation, cryosections of embryos (10 µm) were stained with Congo red, a dye that selectively binds β-sheet-rich amyloid structures and has been used in zebrafish to detect amyloid structures (*33*). In drP18 eGFP and drP18 C50A eGFP expressing embryos, a Congo red-positive signal was detected in a subset of cells and it co-localized with GFP fluorescence, suggesting the presence of amyloid structures in some cells (Fig. 5*C*, top and center). In contrast, eGFP control expressing embryos did not display any Congo red-positive structures under otherwise identical conditions (Fig. 5*C*, bottom). These observations were consistent across biological replicates (n = 8 embryos per condition), and the experiment was independently repeated twice with similar results; representative images are shown. To further probe for the presence of amyloid structures, polarized light microscopy revealed apple-green birefringence in regions corresponding to Congo red staining in drP18 eGFP and drP18 C50A eGFP expressing embryos (Fig. 5*D*), a hallmark feature for the presence of amyloid structures which was missing for the eGFP control.

Together these data suggest that drP18 can be oxidized by endogenously present compounds, leading to assembly into amyloid structures *in vivo*.

## Discussion

Historically, amyloid formation has been associated with intrinsically disordered or pathological proteins, but it is becoming clear that folded, functional proteins can also undergo amyloid transitions under physiological conditions. In these systems, amyloid formation is generally thought to require transient destabilization of the native fold, allowing aggregation-prone conformations to become accessible through local unfolding events, for example, via thermal fluctuations or destabilizing mutations (*3*). Our previous work on human p16 suggested an alternative mechanism where the initiation of amyloid formation is tightly regulated by cellular redox processes, with site-specific cysteine oxidation functioning as a molecular switch that initiates the transition into the amyloid state and cysteine reduction driving disassembly (*22*, *23*). Here we present an example where unlike by simple single-cysteine oxidation, the P18 transition is regulated through a cooperative redox mechanism in which the two cysteine residues fulfil distinct regulatory and executioner functions.

Both cysteines of drP18, C50 and C128, are necessary for a stable, well-folded monomeric protein. Mutation of either or both residues significantly reduces protein stability, as determined by melting temperature measurements. Both cysteine residues are accessible on the protein surface but are separated by a full ankyrin repeat and located on opposite sides of the α-helical motif, with their Cα atoms approximately 21 Å apart. Formation of an intramolecular C50–C128 disulfide bond therefore requires a high degree of flexibility that substantially disrupts the structure of the linear repeats. This intramolecular disulfide-bonded conformation is stable in solution and does not aggregate, it is representing a stable off-state preventing amyloid formation upon oxidation.

There are major structural differences of each homomeric cross-linked species. When only C50 is present, oxidation with most oxidants does not lead to dimerization, but peroxymonocarbonate produces dimeric species that do not further assemble. In contrast, when only C128 is present, oxidation leads to rapid formation of intermolecular dimers and subsequent amyloids. These differences highlight the distinct properties of each cysteine residue in the wild-type protein. C50 is slightly more reactive and oxidizes first, while C128 is the residue that drives intermolecular crosslinking and fibril assembly. NMR spectroscopy highlighted the structural changes at amino acid resolution upon oxidation. Chemical shift perturbations indicated that C128 is significantly affected by C50 oxidation, despite the 21 Å separation, consistent with a long-range regulatory effect of C50 on the C128 molecular environment.

Glutathione is the most abundant intracellular thiol, and the GSH/GSSG system is a cell’s dominant redox buffer, with its ratio shifting in response to different oxidants (*34*) and glutathione can also covalently modify proteins (*35*). In drP18, *S*-glutathionylation of C50 directly controls the ability of C128 to form intermolecular disulfide bonds that trigger the transition into amyloid structures. By blocking C50’s capacity to form the intramolecular disulfide with C128, glutathionylation enables C128 to undergo intermolecular crosslinking upon oxidation. Even in the cysteine-free mutant, glutathione caused detectable chemical shift changes in NMR spectra, suggesting that glutathione interacts with the protein surface independently of covalent modification.

Different oxidants are able to convert the glutathionylated C50 species into amyloid structures, and the resulting amyloid morphology varies between oxidants. This variation likely reflects the rate of dimer formation and different kinetics of amyloid growth. The overall formation rates range from fast with stoichiometric hypothiocyanous acid to slow with hydrogen peroxide. The type of oxidant may therefore act as a regulator of polymorphism, with potential functional consequences. HOSCN is an immune system-derived oxidant formed by enzymes such as myeloperoxidase, which has been reported in zebrafish (27), however, it is unclear if thiocyanate is a relevant metabolite in this organism.

Another major feature of drP18 amyloid structures is their reversibility. Upon addition of reducing agents, the glutathionylated drP18 variant showed disappearance of dimeric species by SDS-PAGE and a decrease in ThT fluorescence, consistent with disulfide-mediated disassembly. In contrast, a fraction of the C50S mutant remained dimeric, and ThT fluorescence decreased significantly only for peroxymonocarbonate-treated samples. C50 therefore appears to contribute to the disassembly pathway as well, either through a direct effect of the cysteine side chain or through conformational effects on intermediates. These findings further support the conclusion that different oxidants produce amyloid morphologies with distinct properties.

When overexpressed in zebrafish embryos, drP18 formed aggregates displaying amyloid-type features and disulfide crosslinks, potentially triggered by endogenous oxidants and supporting a role for redox-controlled amyloid formation *in vivo*.

Together, these data reveal a multi-step redox mechanism that allows drP18 to switch between two major structural states. The reversible transition from a monomeric, α-helical conformation into β-sheet-based amyloid structures involves a major structural rearrangement, including the hydrogen bond network that stabilizes each state. The ability of drP18 to inhibit CDK4 is present in its monomeric form, including under glutathionylated conditions, but is lost upon conversion into amyloid assemblies and restored upon reduction.

Cysteine redox switches have been reported for proteins involved in vital cellular processes, and reactive oxygen species are recognized as important second messengers in parallel to their deleterious effects under conditions of high oxidative stress (*36–38*). One prominent example is PTEN, where oxidation of the catalytic cysteine residue induces formation of a reversible intramolecular disulfide bond with a cysteine residue originally located 6 Å apart, thereby inhibiting its phosphatase activity (*39*). Other examples include peroxiredoxins, which are multi-domain scavengers of peroxides with a highly reactive cysteine residue in their active site. These are reduced by a resolving cysteine residue, which in some cases is about 14 Å apart and involves a local unfolding event for their action (*40*). Larger cysteine-cysteine distances have been reported but involve major domain rearrangements (*41*) or intrinsically disordered regions (*42*).

The two cysteine residues in drP18 are about 21 Å apart, the largest separation among these examples, and their oxidation can lead to a dramatic structural rearrangement into amyloid fibrils that is reversible through disulfide bond reduction. This demonstrates that redox switches built on distant cysteine pairs can drive outcomes ranging from local unfolding into less structured species to large-scale structural reorganization, with important functional impact.

## Materials and Methods

### Protein expression and purification

An *Escherichia coli* codon-optimized gene of drP18 wild-type (Uniprot ID: B0UXZ0) and cysteine mutants were generated by Genscript Biotech and each cloned separately into a pETZ2 vector with an N-terminal 6× histidine tag followed by a Z-tag (protein A) and a Tobacco etch virus (TEV) protease cleavage site, leaving a glycine and an alanine residue at the N-terminus. Bacterial transformation was carried out using *E. coli* BL21 (DE3) competent cells and a selected colony was inoculated in the autoclaved LB media containing Kanamycin antibiotic (50 μg/L). After overnight incubation at 28 °C, glycerol stocks were prepared with 1:1 ratio of concentrated cell culture to glycerol (% v/v) and stored at -80 °C. Uniform ^15^N and/or ^13^C uniformly isotopic labeling of proteins was accomplished by culturing cells in minimal medium in which ^15^NH₄Cl and/or ^13^C-glucose served as the sole nitrogen and carbon sources. Protein synthesis was initiated with 0.5 mM of isopropyl-1-thio-D-galactopyranoside (IPTG) once the OD_600_ of the bacterial culture was approximately 0.4-0.6. The cells were harvested by centrifugation and resuspended in purification buffer (440 mM potassium acetate, 80 mM HEPES [4-(2-hydroxyethyl)-1-piperazineethanesulfonic acid], 80 mM imidazole, 20% v/v glycerol, pH 8.0) containing freshly added 2 mM BME (β-mercaptoethanol). The cell suspension was then ultrasonicated for lysis and the supernatant was applied to a pre-equilibrated Ni-NTA agarose column (Qiagen). After washing the suspension with purification buffer, the protein was eluted using purification buffer containing 200 mM imidazole. Size exclusion chromatography of the eluate was then performed using a HiLoad 16/600 Superdex 75 pg column (Cytiva) on an Äkta purification system. The fractions containing the protein were pooled and incubated with TEV protease (∽1:50 molar ratio) overnight at 4 °C. Next day, the protein solution was again applied to the Ni-NTA agarose column to remove the tagged cleavage products, 6× histidine-tagged TEV and undigested protein. The cleaved protein fraction was treated with 2 mM tris(2-carboxyethyl)phosphine (TCEP) for 15 min at 4 °C and buffer exchanged with 4 mM HEPES containing 200 μM ethylenediaminetetraacetic acid (EDTA) (pH 7.4) on a HiPrep 26/10 desalting column. The protein was then concentrated to 70 µM and stored at -80 °C until further use.

### SDS-PAGE analysis

Diamide, hydrogen peroxide (H_2_O_2_), peroxymonocarbonate (HCO_4_ ^-^), and hypothiocyanous acid (HOSCN) were prepared as previously described (*29*). Following oxidant treatment, protein samples were treated with 10 mM NEM (N-ethylmaleimide) to block the remaining free cysteine side chain sites. Protein samples were prepared in SDS-PAGE loading buffer by diluting the 4× stock to a final 1× concentration, with or without the addition of the reducing agent dithiothreitol (DTT) and resolved using Bolt 4-12% Bis-Tris Plus (Invitrogen) gels at 200 V. Molecular weights were estimated by reference to Precision Plus Protein Dual Color Standards (Bio-Rad).

### Thioflavin T fluorescence assay

Protein samples (20 μM) were mixed with 10 μM Thioflavin T (ThT) dye in a 96 well-plate (Corning CLS3881) covered with a MicroAmp optical adhesive film (Thermo Fisher Scientific) and adjusted to 25 μL with 4 mM HEPES (pH 7.4). The oxidant was added right before the start of the assay. The assay was run over a time course of 24 hours or longer at 25 °C and fluorescence was measured at 2-minute intervals with emission and excitation wavelengths of 482 nm and 435 nm, respectively, using a Molecular Devices M5 microplate reader. Data was normalized using Amylofit and half time of amyloid formation was calculated through its inbuilt function (*43*).

### Differential Scanning Fluorimetry

From a 5000× stock solution, 10× concentration of the SYPRO-orange dye was added to a purified protein sample (20 μM) and the final volume of 30 μL was generated by using 4 mM of HEPES buffer (pH 7.4). Samples were pipetted in a 96 well-plate (Corning CLS3881) covered with a MicroAmp optical adhesive film (Thermo FisherScientific). All DSF measurements were performed in quintuplicates on a QuantStudio 3 real time PCR machine where the temperature of the plate was gradually increased from 10 °C to 95 °C at the rate of 0.015 °C per second and the results obtained were analyzed using the Protein Thermal Shift Software 1.4.

### Solution NMR spectroscopy

500 μL of ^15^N-labelled protein samples were prepared in a 4 mM HEPES buffer (pH 7.4) containing 200 μM EDTA and 50 μL D₂O was added to each sample as a lock solvent. Two-dimensional (^1^H^15^N) HSQC (Heteronuclear single quantum coherence spectroscopy) spectra were acquired on a Bruker 600 MHz spectrometer operated by an Avance III console equipped with a TXI triple-resonance probe including z-gradients. The sample temperature was maintained at 298 K throughout the measurement. The acquired data was processed and analyzed using Topspin 4.2.0. The exported data was further visualized and analyzed using CCPNMR analysis 3.2.0 (*44*).

To determine the midpoint redox potential (E₀) for protein *S*-glutathionylation, redox titrations were performed in the presence of 4 mM reduced glutathione (GSH), while the concentration of oxidized glutathione (GSSG) was systematically increased (0.05, 0.10, 0.20, 0.40, 0.80, 1.30, 2.30, 3.90, 5.90, 7.80, 8.80, 9.80, 11.70, 13.00, and 15.00 mM). After each addition, the sample was gently mixed, and an HSQC spectrum was recorded immediately. Standard ¹H¹⁵N HSQC spectra were recorded using the same settings as the reference scan, with the number of scans increased as needed (initially n = 12, then 20, 24 and 32) to maintain comparable signal-to-noise ratios as the sample volume increased. The redox potential of the GSSG/2GSH couple at each titration point was calculated using the Nernst equation, and the data were analyzed and fitted using GraphPad Prism, as described previously (*22*).

### Atomic Force Microscopy (AFM)

The protein sample was oxidized at 25 °C and applied to a charged mica surface. The mica was functionalized by cleaving to expose a fresh surface and incubating with 0.05% (v/v) (3-aminopropyl)triethoxysilane (APTES) in Milli-Q water for 10 min. Excess APTES was removed by rinsing thoroughly with Milli-Q water, and the surface was gently dried using compressed nitrogen gas. The protein sample was then applied to the functionalized mica surface and incubated for a further 10 min. Unbound protein was removed by rinsing with Milli-Q water, and the mica was again gently dried under compressed nitrogen. Samples were imaged using a Bruker Dimension Icon-IR atomic force microscope with a spatial resolution of 10 nm. AFM images were processed and analyzed using NanoScope Analysis software (version 3.0).

### Negative-stain transmission electron microscopy

drP18 samples (20 μM) were oxidized for 24 h at 25°C with the different oxidants. Carbon-coated copper grids (ProSciTech) were floated on a 7 μL droplet of the oxidized sample for 60 s, blotted to remove excess solution, and floated on water for 30 s before being blotted again. Grids were then stained with 2% (w/v) uranyl acetate by floating on a 7 μL droplet for 30 s, followed by blotting of excess stain with filter paper. Grids were dried overnight and visualized using a ThermoScientific Talos L120C transmission electron microscope. Multiple grid meshes were examined for each sample.

### Kinase assay

Proteins were oxidized for 24 h with equimolar HOSCN or with a 10-fold protein-to-oxidant ratio for diamide and peroxymonocarbonate, followed by isolation of the amyloid fraction using a Superdex 200 Increase 10/300 column (Cytiva, Marlborough, MA, USA) connected to an ÄKTA pure protein chromatography system. For reduction experiments, the isolated amyloid fraction was concentrated and the protein concentration was determined by NanoDrop. Samples were then incubated with a 10-fold molar excess of DTT relative to the oxidant concentration at 25°C for 24 h. The samples were subsequently buffer exchanged (HiPrep 26/10, Cytiva) to remove excess DTT before re-oxidation with the indicated oxidant.

Oxidized, reduced, and unoxidized protein samples (1.05 μM) were added to 350 nM human GST-tagged CDK4/Cyclin D1 complex (Invitrogen, Carlsbad, CA, USA, Cat. No. PV4400). 6×His– Rb substrate (16.5 kDa; 3.5 μM; MyBioSource, Cat. No. MBS143254) and ATP (100 μM) were added to the reaction mixture. Reactions were performed in kinase buffer (125 mM Tris-HCl, pH 7.5, 50 mM MgCl₂) at 28°C for 30 min and terminated by boiling at 95°C for 10 min in 4× reducing SDS buffer. Rb phosphorylation was assessed by immunoblotting using an anti-pRB antibody (Cell Signaling Technology).

### Zebrafish maintenance and breeding, genomic DNA extractions and embryo microinjections

Adult zebrafish and their embryos were maintained at 27–28°C at the Otago Zebrafish Facility (Department of Pathology and Molecular Medicine, University of Otago, Dunedin, New Zealand). All animal experiments were conducted where appropriate under the approval of the University of Otago Animal Ethics Committee (SOP-OZF-004, AUP 25-72). Adult WIK (wild-type) zebrafish were paired and placed in breeding tanks one day prior to breeding with a temporary divider placed in between males and females. The following day, the divider was removed to enable natural spawning. The eggs were collected using a mesh strainer and transferred into a petri dish with embryo medium (E3). For microinjection, embryos were collected specifically at the one-cell stage and injected under a dissection microscope with 200 pg of the indicated mRNA constructs using a fine borosilicate capillary needle. Following injection, embryos were maintained at 28 °C in E3 and allowed to develop to 24 hpf prior to collection and processing for downstream analyses.

Zebrafish adult fin clips were digested in 50 µL of 50 mM NaOH at 95 °C for 20 minutes. This lysis reaction was then neutralized with 5 µL 1M Tris-HCl (pH 8.0). Genomic DNA was stored at -20 °C until required for Sanger sequencing.

### Zebrafish embryo processing, western blot analysis, and Congo red staining

For western blot analysis, microinjected embryos were pooled in groups of 30 (n = 30), manually dechorionated, and transferred to ice-cold Ringer’s solution containing 1 mM EDTA (pH 7.0) and protease inhibitors. Embryos were washed twice in fresh cold Ringer’s solution and pelleted by centrifugation for 1–2 min. Excess liquid was removed and samples were resuspended in 150– 200 µl non-reducing SDS sample buffer (0.63 ml 1 M Tris-HCl, pH 6.8, 1.0 ml glycerol, 1.75 ml 20% SDS, and 6.12 ml H₂O; total volume 10 ml). Samples were boiled for 5 min, centrifuged for 1–2 min, and analyzed by SDS-PAGE and western blotting.

For Congo red staining, embryos were fixed overnight in 4% paraformaldehyde at 4 °C, cryoprotected in sucrose, embedded in OCT compound, and cryosectioned at 10 µm thickness. Sections were mounted onto Trajan Series 1 adhesive microscope slides and allowed to adhere overnight at room temperature. Slides were stained with 0.5% Congo red prepared in 50% ethanol for 20 min, washed three times in Milli-Q water followed by a final rinse in PBS, and subsequently incubated with DAPI (0.5 µg/ml in PBS) for 10 min. Sections were rinsed in PBS, mounted, and visualized using a Zeiss fluorescence microscope.

### Preparation of amyloid fibrils

To generate amyloid fibrils, glutathionylated drP18 or the drP18 C50S mutant (20 or 40 µM) was incubated with diamide for 24 or 48 h. Following oxidation, excess reagent was removed by buffer exchange into Milli-Q water using a HiTrap™ Desalting 5 mL column (Cytiva) on an ÄKTA chromatography system. The desalted samples were then incubated at 28 °C until visible precipitates formed (typically within 24 h after buffer exchange). Samples were centrifuged at 21,000 × g for 1 h at 4 °C to pellet the fibrillar material. The pellet was washed once with Milli-Q water, centrifuged again under the same conditions, and finally resuspended in a minimal volume of Milli-Q water for storage at 4 °C until further analysis.

### Whole protein mass spectrometry

Whole protein samples were analyzed using a Velos Pro ion trap mass spectrometer, coupled to a Dionex UltiMate 3000 HPLC system, with a 50 µL injection loop (Thermo Fisher Scientific, Waltham, MA, USA). Samples were stored on the autosampler tray at 5 °C. An Accucore-15-C4 HPLC column (50 mm × 2.1 mm, 2.6 μm; Thermo Fisher Scientific, Waltham, MA, USA) was used for chromatographic separation using 100% water (0.1% formic acid) as solvent A and 100% acetonitrile (0.1% formic acid) as solvent B. The column temperature was set to 60 °C. The column was equilibrated with 90% solvent A and 10% solvent B for 2.5 min, then a linear gradient was run for 2.1 min to 20% solvent A and 80% solvent B to elute the proteins. The column was then flushed with 20% solvent A and 80% solvent B for 2.5 min, then re-equilibrated at initial conditions for 2.5 min. A flow rate of 0.4 mL/min was used, and approximately 1 µg of protein was injected for each sample. Nitrogen was used as sheath gas. The temperature of the heated capillary was 275 °C. Mass spectra (400–2000 m/z) were averaged over the full length of each protein peak using Thermo Scientific Xcalibur 4.2.4.7 (Thermo Fisher Scientific, Waltham, MA) and deconvoluted to yield the molecular masses using ProMass for Xcalibur 2.8.5 (Novatia LLC, Monmouth Junction, NJ, USA).

### Tryptic digest mass spectrometry

Approximately 10 μg of protein was oxidized for 24 h with the desired oxidant followed by NEM (10 mM) addition. A 50:1 substrate:trypsin protease (Promega) weight ratio was used to perform digestion of the samples, with overnight incubation at 25°C.

The digested protein samples were then injected onto a Jupiter 4 μm Proteo 90 Å column with the dimensions of 150 x 2.0 mm (Phenomenex, Torrance, CA, USA) and the column oven was set to a temperature of 40 °C. The column was equilibrated with 95% Solvent A and 5% Solvent B for 5 min before a linear gradient was run for 20 min to 55% Solvent A and 45% Solvent B in order to achieve separation. The column was then flushed with 5% Solvent A and 95% Solvent B for 5 min and subsequently re-equilibrated in the initial conditions for 5 min. A flow rate of 0.2 mL/min was used throughout experiments.

The eluted tryptic peptides were analyzed using a data-dependent nth order double play scan procedure. The ten most abundant peaks in a full scan (300–2000 m/z) were sequentially selected, and an MS/MS scan was performed using collision induced dissociation with a normalized collision energy of 40%. MS/MS spectra were recorded using dynamic exclusion with a repeat count of 3, a repeat duration of 30 seconds, and an exclusion duration of 60 seconds. For quantification of the tryptic peptides, peak areas were obtained by filtering the acquired mass spectra with the predicted m/z values of the expected charge states of each of the tryptic peptides (within the 300–2000 m/z range) and using Genesis peak detection (Gaussian 7-point smoothing). The identity of each of the detected peptides was confirmed by comparing the acquired corresponding MS/MS fragmentation patterns to the theoretical peptide fragment ions predicted using mMass software (version 2.4).

For quantification of disulfide-linked tryptic peptides, the m/z values of peptides of interest were predicted and collision-induced dissociation-MS/MS spectra were acquired for each of them in positive-ion mode. The collision energy was set at 40. Peptide fragments were assigned by comparing the acquired corresponding MS/MS fragmentation patterns to the theoretical peptide fragment ions predicted using mMass software. Each peptide species was quantified by post-acquisition filtering the MS/MS spectra obtained for a chosen abundant and characteristic fragment ion and then measuring the area under the curve of the resulting peak (peak algorithm: Genesis, peak smoothing: Gaussian 7 points).

## Acknowledgements

We thank Thomas Evans for assistance with atomic force microscopy measurements, Duane Harland and Marina Richena for assistance with electron microscopy measurements and Hector Ulises Mancilla Diaz for assistance with polarized light microscopy. We thank Michael Meier for providing zebrafish DNA samples. We further thank Tony Kettle and Christine Winterbourn for highly valuable input, Nina Dickerhof for support with mass spectrometry experiments and the members of the Mātai Hāora - Centre for Redox Biology and Medicine for stimulating discussions.

## Author contributions

Conceptualization: C. G., V. K. M., A. S. and H. D. Methodology: A. S., H. D. and C. G. Formal analysis: A. S., H. D., N. J. M., G. G., K. M. O., A. D. B., P. C., R. C. and J. H. Writing—original draft: A. S. and C. G. Writing—review and editing: A. S., H. D., V. K. M. and C. G. Visualization: A. S. Supervision: C. G., V. K. M., H. D. and J. H. Project administration: C. G., A. S. and H. D. Funding acquisition: C. G. and V. K. M. The authors declare that they have no competing interests. All data needed to evaluate the conclusions in the paper are present in the paper and/or the Supplementary Materials.

