## Supplementary for "Hierarchical cysteine oxidation controls reversible amyloid formation in an ankyrin repeat protein"

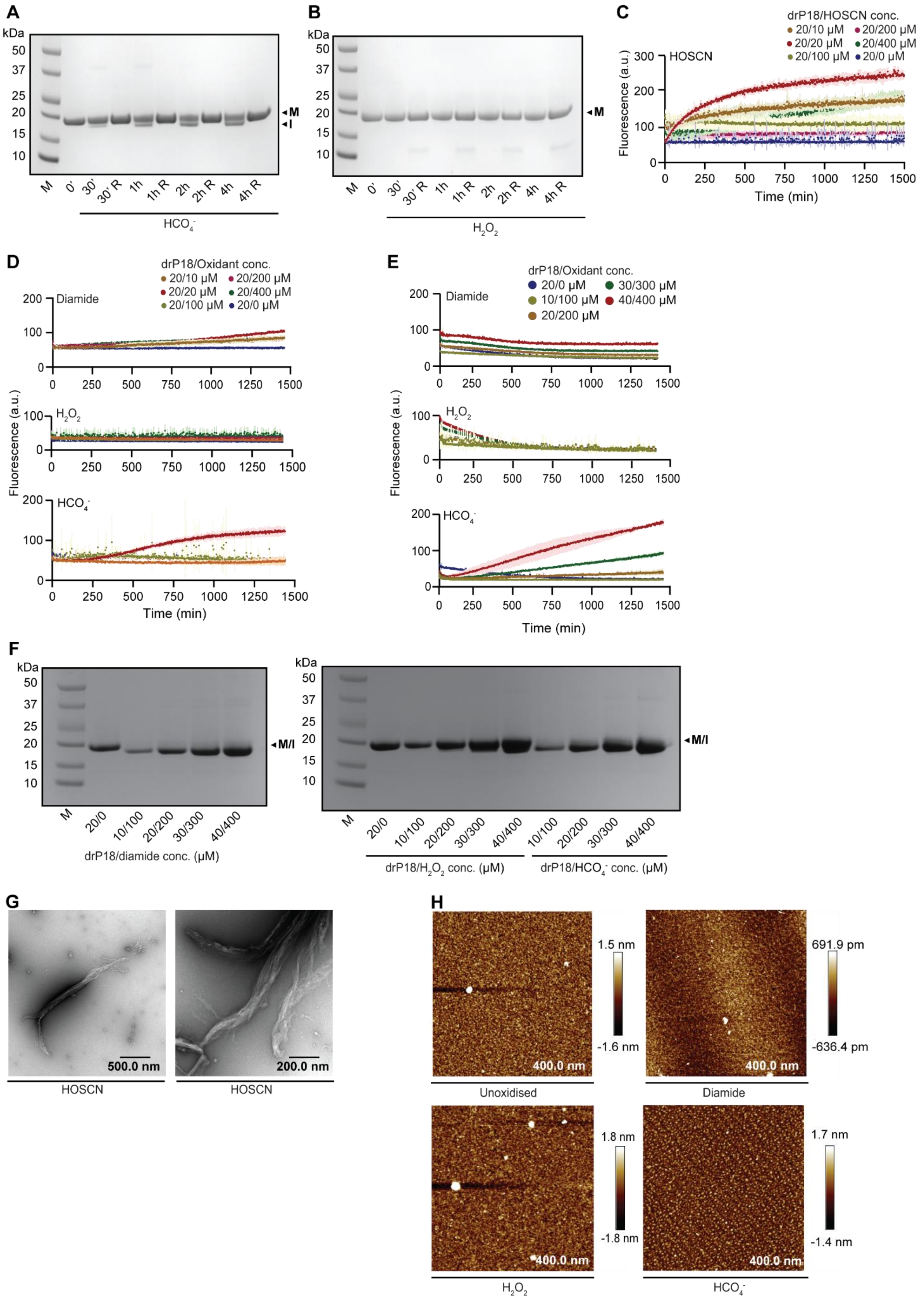
A
kDa
50
37
25
20
15
10
M
I
M
0'
30'
30' R
1h
1h R
2h
2h R
4h
4h R
HCO4-
B
kDa
50
37
25
20
15
10
M
M
0'
30'
30' R
1h
1h R
2h
2h R
4h
4h R
H2O2
C
drP18/HOSCN conc.
20/10 μM
20/200 μM
20/20 μM
20/400 μM
20/100 μM
20/0 μM
HOSCN
Fluorescence (a.u.)
Time (min)
D
drP18/Oxidant conc.
20/10 μM
20/200 μM
20/20 μM
20/400 μM
20/100 μM
20/0 μM
Diamide
H2O2
HCO4-
Fluorescence (a.u.)
Time (min)
E
drP18/Oxidant conc.
20/0 μM
30/300 μM
10/100 μM
40/400 μM
20/200 μM
Diamide
H2O2
HCO4-
Fluorescence (a.u.)
Time (min)
F
kDa
M/I
M
20/0
10/100
20/200
30/300
40/400
drP18/diamide conc. (μM)
drP18/H2O2 conc. (μM)
drP18/HCO4- conc. (μM)
G
500.0 nm
200.0 nm
HOSCN
HOSCN
H
1.5 nm
-1.6 nm
400.0 nm
Unoxidised
691.9 pm
-636.4 pm
400.0 nm
Diamide
1.8 nm
-1.8 nm
400.0 nm
H2O2
1.7 nm
-1.4 nm
400.0 nm
HCO4-

**Figure S1. Oxidation of drP18 with different oxidants.** (A) SDS-PAGE analysis of 20 µM drP18 (M) shows formation of I, an intramolecular disulfide bond following oxidation with 200 µM $HCO_4^-$ over a time course of 4 h. Addition of the reducing agent DTT (R) breaks the disulfide bond. (B) SDS-PAGE analysis of 20 µM drP18 (M) following oxidation with 200 µM $H_2O_2$ over a time course of 4 h. The protein predominantly remains monomeric. (C) Thioflavin-T fluorescence kinetics assays of drP18 (20 µM) oxidized with HOSCN at concentrations ranging from 10–400 µM. (D) Thioflavin-T fluorescence kinetics assays of drP18 (20 µM) oxidized with different oxidants at concentrations ranging from 10–400 µM. (E) Thioflavin-T fluorescence kinetics assay of drP18 at varying protein concentrations (10–40 µM), oxidized with different oxidant concentrations (10–400 µM) while maintaining a 1:10 protein-to-oxidant ratio. (F) SDS-PAGE analysis of drP18 protein (M) across a concentration series at a 1:10 protein-to-oxidant ratio at 25 °C shows formation of I, an intramolecular disulfide bond. (G) Representative electron microscopy (EM) images of 20 µM drP18 assemblies formed when treated with 20 µM HOSCN for 24 h. Scale bar: 500 nm. (H) Representative AFM images of 20 µM drP18 in its unoxidized state (left) and after oxidation with 200 µM diamide (center left), $H_2O_2$ (center right) and $HCO_4^-$ (right) for 24 h. Scale bar: 400 nm.

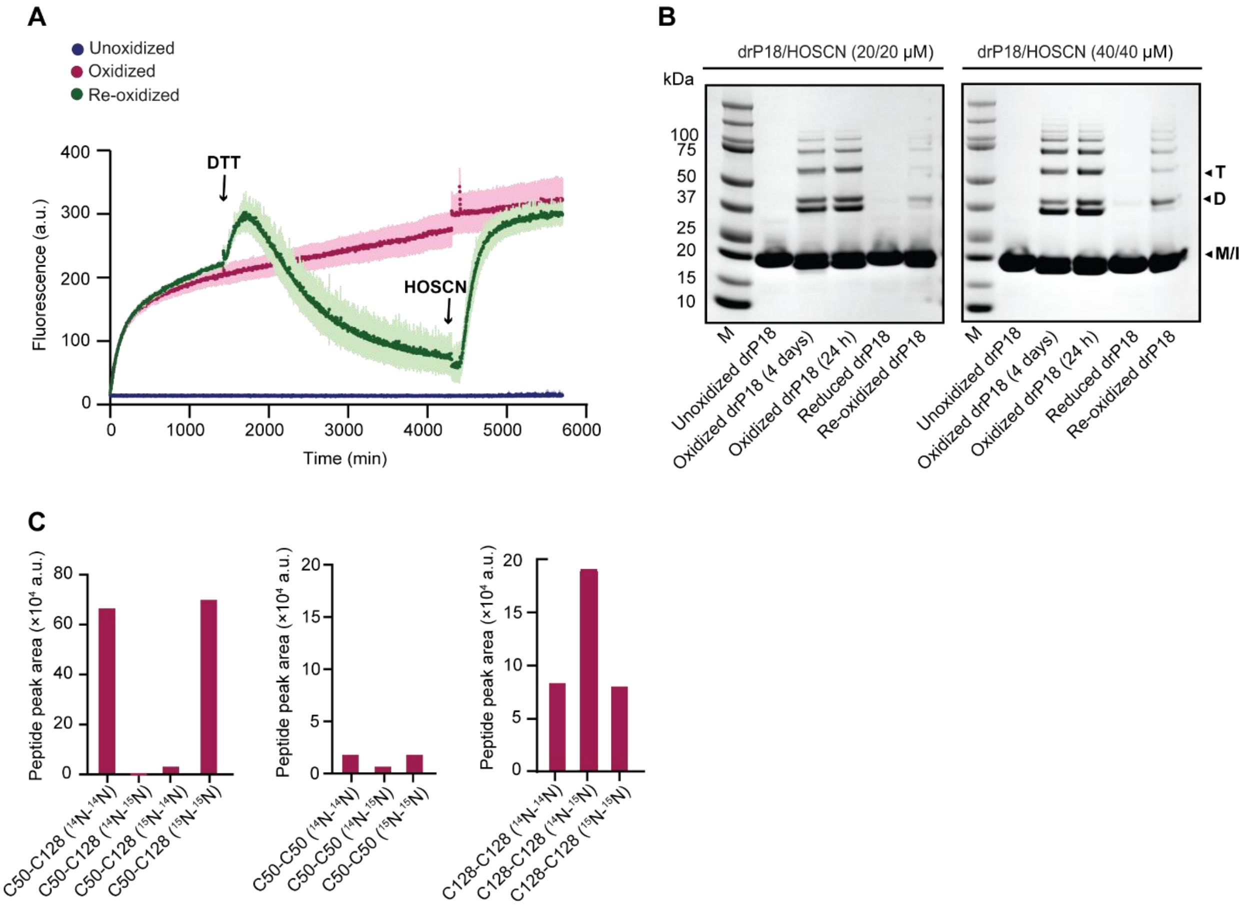


**Figure S2. Oxidation of drP18 and mass spectrometry analysis using different conditions.** (A) Thioflavin-T fluorescence kinetics assay of drP18 (40 µM) oxidized with HOSCN (40 µM). Addition of 400 µM DTT after 1400 minutes (green curve and addition indicated by arrow) causes reduction of fluorescence to baseline and subsequent re-addition of 840 µM HOSCN at 4400 minutes (excess to quench residual DTT) increases the ThT fluorescence. The control curves (pink, oxidized and blue, unoxidized) behave as described before. (B) SDS-PAGE analysis of drP18 (20 µM, left and 40 µM, right) oxidized with HOSCN (20 µM and 40 µM, respectively). Addition of 200 and 400 µM DTT (left and right, respectively) after 24 h reduces the disulfide bond and subsequent re-addition of 420 µM and 840 µM HOSCN shows re-formation of D, potential dimer and T, potential trimer. (C) LC-MS/MS analysis of disulfide-linked peptides following oxidation of mixed 14N- and 15N-labelled drP18 with a 1× molar equivalent of HOSCN. Equal amounts of 14N- and 15N-labelled protein were mixed prior to oxidation, free thiols were alkylated with *N*-ethylmaleimide (NEM), and proteins were digested with trypsin before analysis. Intramolecular disulfides generate only $^{14}$N-$^{14}$N and $^{15}$N-$^{15}$N peptide species, whereas intermolecular disulfides produce the characteristic $^{14}$N-$^{14}$N:$^{14}$N-$^{15}$N/$^{15}$N-$^{14}$N:$^{15}$N-$^{15}$N isotopic pattern (1:2:1). C50-C128 disulfide showing only homoisotopic peptide species, consistent with intramolecular disulfide formation (left). C50-C50 disulfide, with only low, inconsistent signal (center). C128-C128 disulfide showing the expected 1:2:1 isotopic distribution, consistent with intermolecular disulfide formation (right).

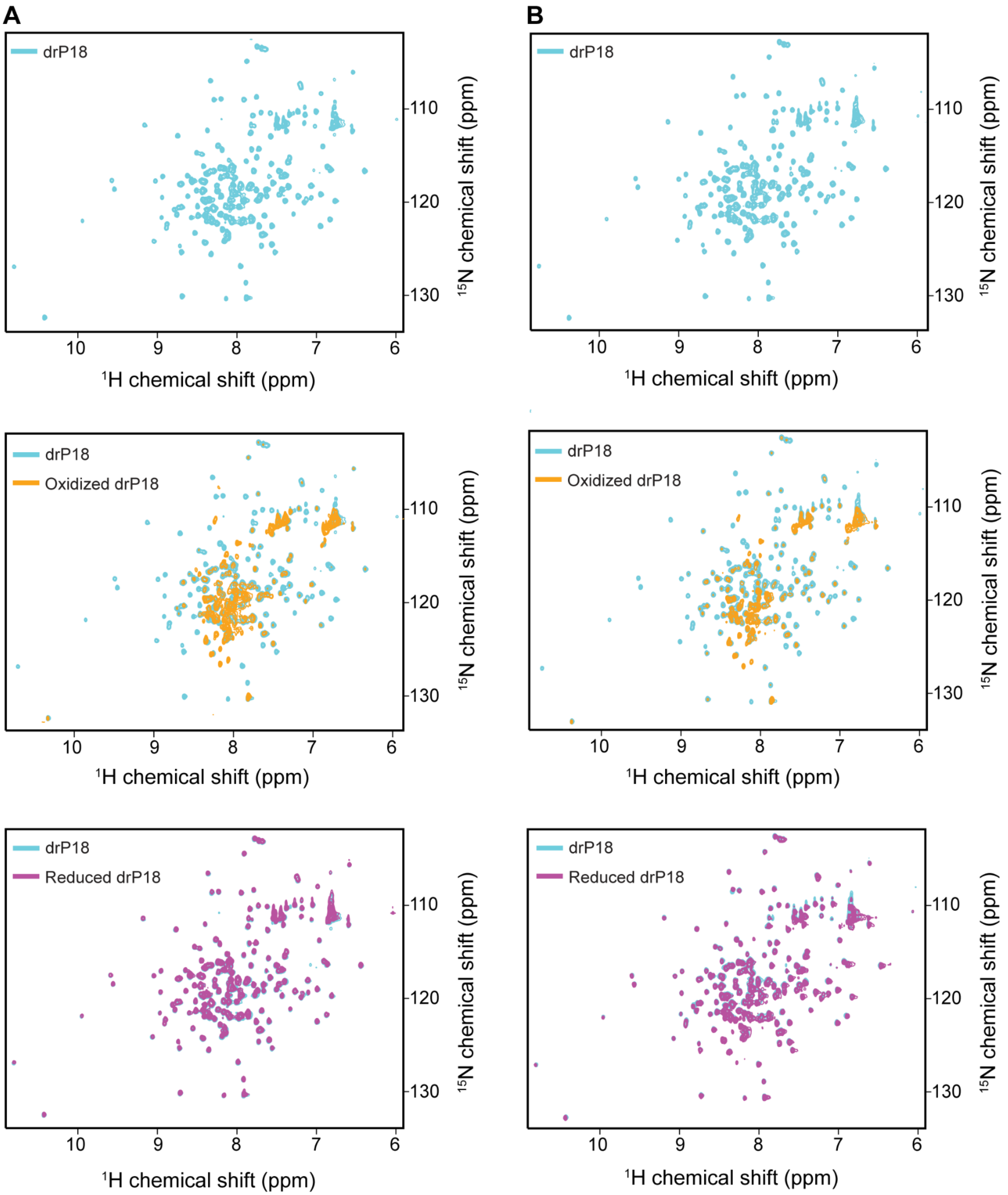


**Figure S3. Oxidation of drP18 and analysis by solution NMR spectroscopy.** Two-dimensional $^{1}H^{15}N$ HSQC solution NMR spectra of a uniformly $^{15}N$ labelled, 170 μM sample of drP18 at 600 MHz and 298 K. A reference HSQC spectrum of drP18 is shown (top, cyan). (A)

and (B) correspond to oxidation with diamide and HOSCN, respectively. For this experiment, diamide (1.7 mM) or HOSCN (170 µM) was added to drP18, and $^{1}H^{15}N$ HSQC spectra were acquired for 24 h incubation at 25 °C. Overlay of HSQC spectra showing oxidized drP18 (middle, orange) compared with the reference spectrum (middle, cyan). Addition of DTT restores the drP18 structure (bottom, magenta). The HSQC spectrum following DTT treatment shows near-complete overlap with native protein, indicating reversibility of the oxidation-induced structural changes.

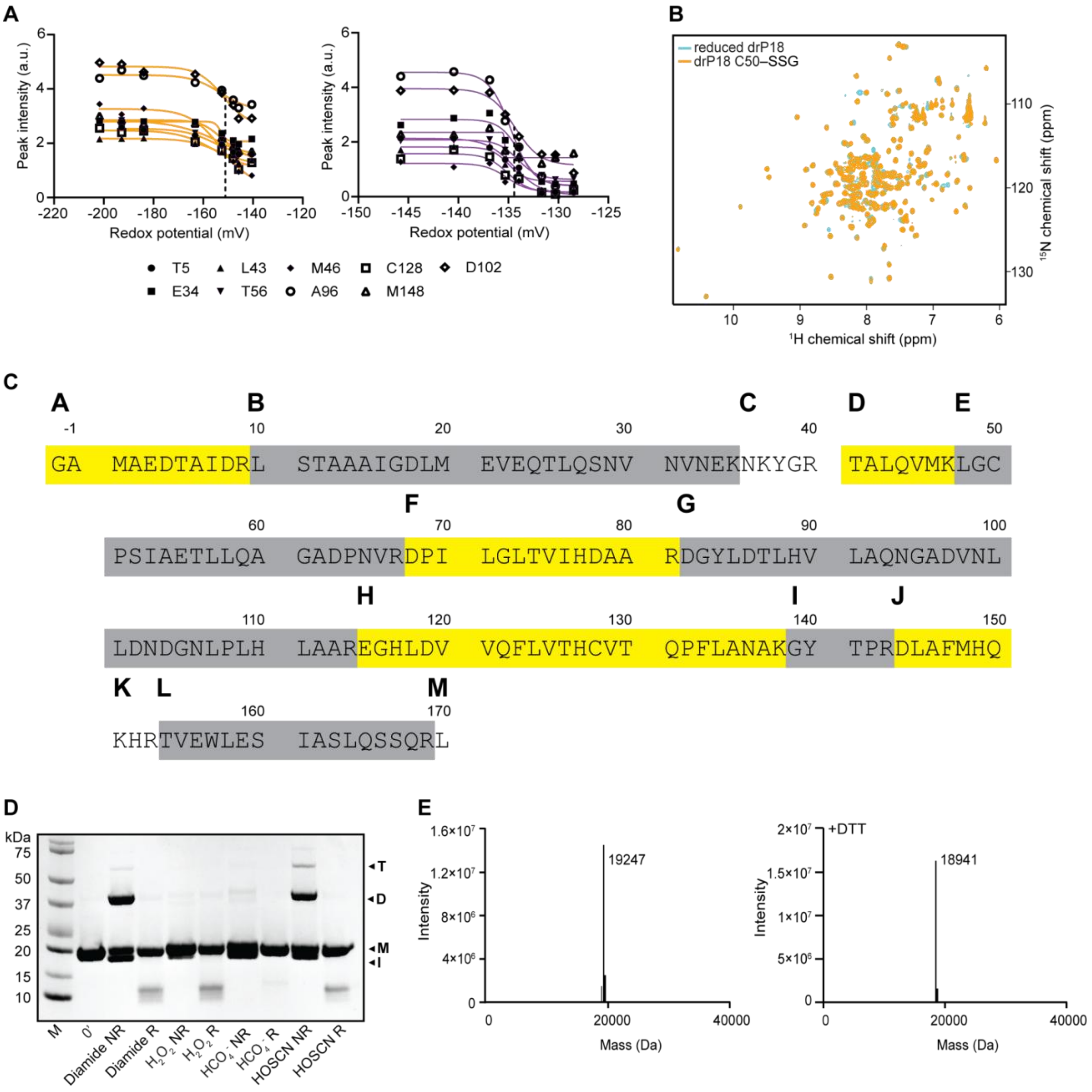


**Figure S4. NMR and mass spectrometry analysis of glutathionylated drP18.** (A) Residues impacted by the redox status of both C50 (orange) and C128 (magenta), with the assignment legend displayed below the graph. (B) $^{1}H^{15}N$ HSQC solution NMR spectrum of drP18 in the reduced state (cyan) and after *S*-glutathionylation of C50 (orange), during the titration series with 0.4 mM GSH and 10 mM GSSG. (C) Full tryptic digestion of drP18 and analysis by mass spectrometry. Depiction of theoretical peptides upon full trypsin digestion (sequentially labeled A–M). Experimentally detected tryptic peptides are highlighted in alternating yellow and gray to illustrate sequence coverage, regions without highlighting have not been observed. (D) SDS-PAGE analysis of 20 µM drP18 C50–SSG (monomeric, M) shows the formation of D, dimers, T, trimers, and I, an intramolecular disulfide-bonded species following oxidation with 200 μM diamide, $H_2O_2$, and $HCO_4^-$, and 20 μM HOSCN over a 4 h time course at 25 °C. DTT addition reverses the disulfide-linked bands. (E) Intact-protein mass spectrometry analysis of drP18 after glutathionylation. The mass spectrum of glutathionylated $^{15}N$-labelled drP18 is showing the single glutathionylated species (expected MW of 19,245.5 Da, detected MW of 19247 Da, left). The mass spectrum following DTT treatment demonstrates loss of the glutathione adduct and restoration of the unmodified protein, consistent with reversible glutathionylation at C50 (expected MW of 18940.4 Da, detected MW of 18941 Da, right).

A

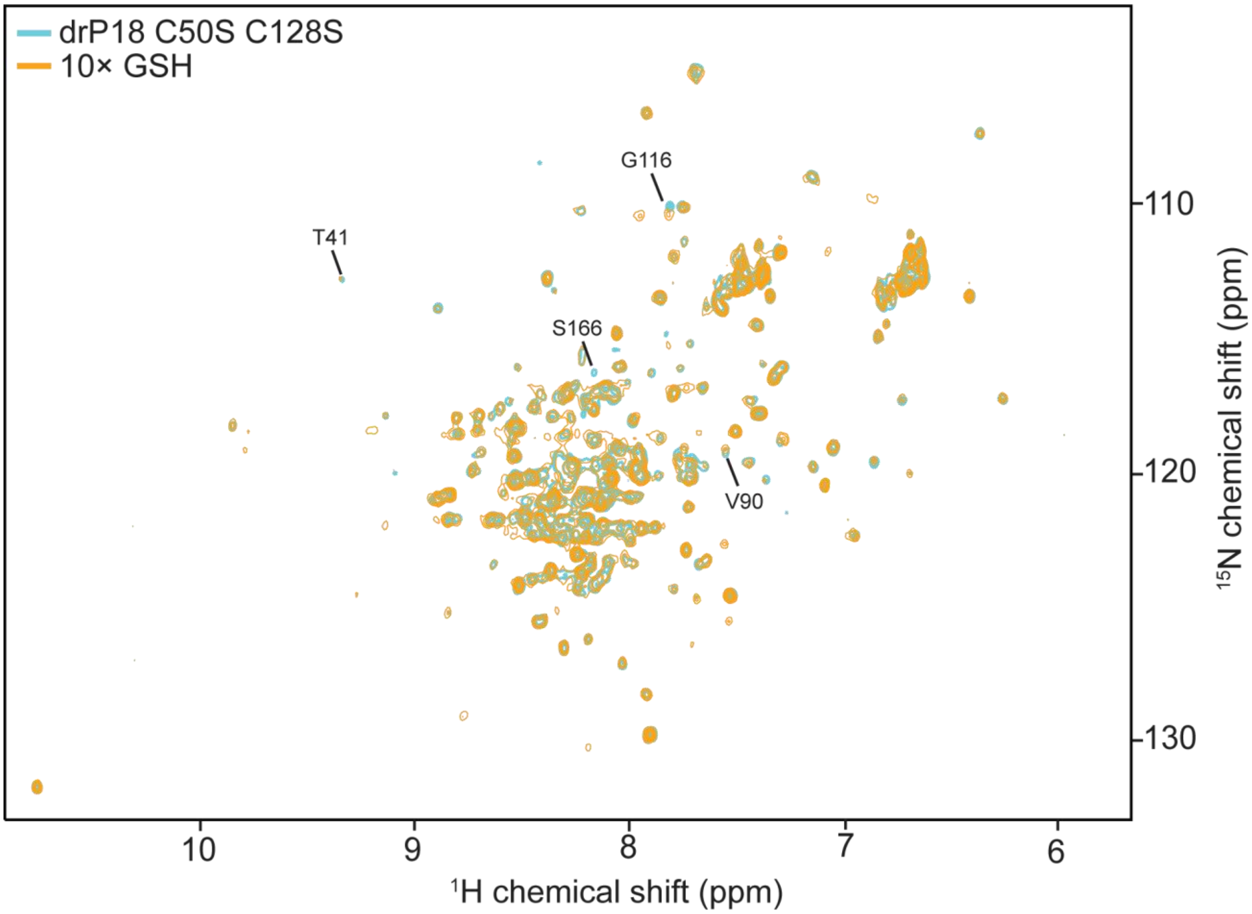


**Figure S5. Two-dimensional $^{1}H^{15}N$ HSQC solution NMR spectra of a uniformly $^{15}N$-labelled drP18 cysteine-free double mutant recorded at 600 MHz and 298 K**. A reference HSQC spectrum of drP18 double mutant is shown (cyan). The HSQC spectrum following addition of 10-fold molar excess GSH is overlaid (orange) with the reference spectrum. Some resonances exhibiting chemical shift perturbations upon GSH addition are highlighted in black. The observed spectral changes indicate weak interaction between GSH and drP18 in the absence of cysteine residues.

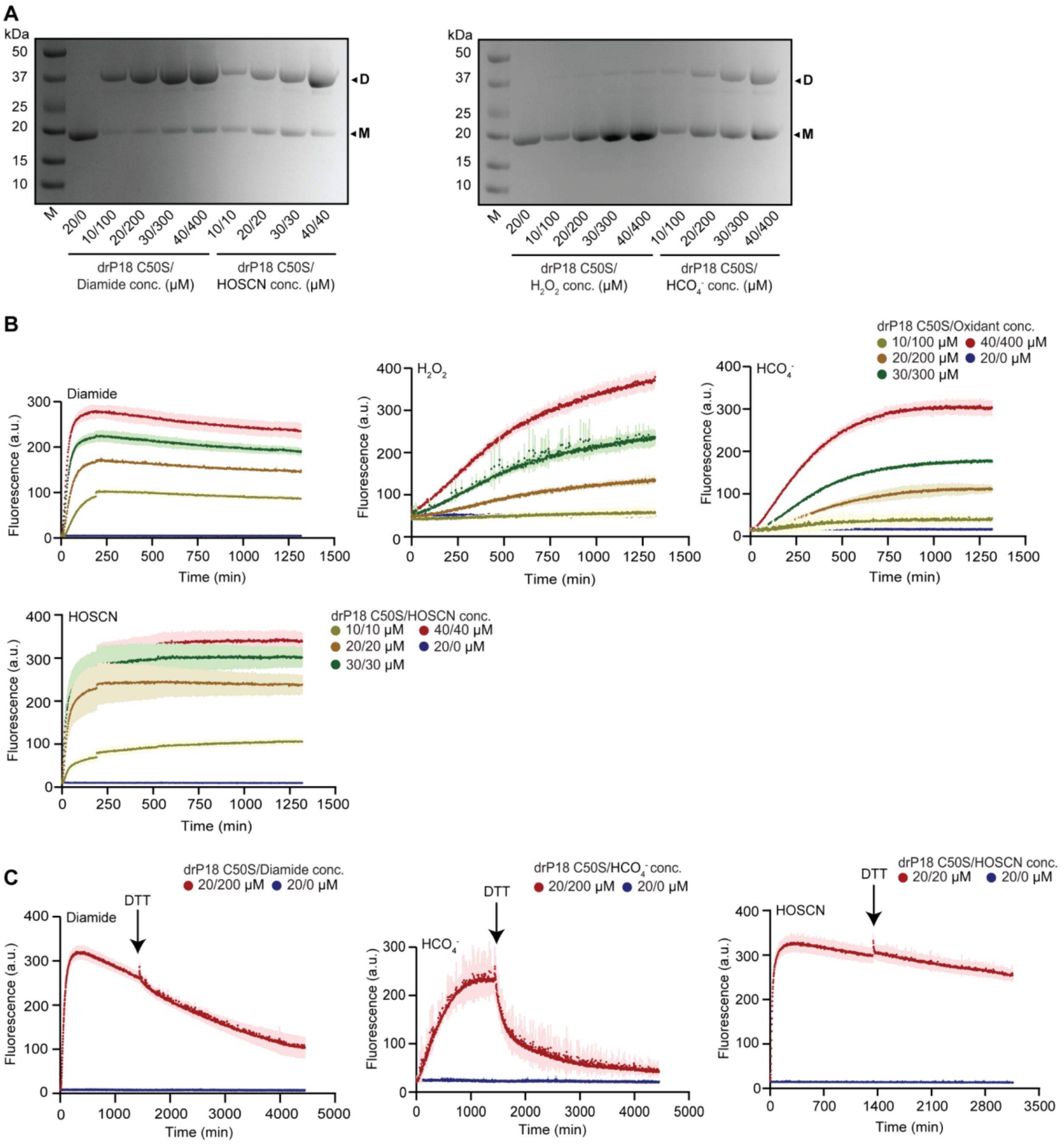


**Figure S6. Oxidation of drP18 C50S with different oxidants.** (A) SDS-PAGE analysis of drP18 C50S protein (M) across a concentration series at a 1:10 protein-to-oxidant ratio at 25 °C for 4 h shows formation of D, dimer, whereas HOSCN was used at a 1:1 protein-to-oxidant ratio. (B) Thioflavin-T fluorescence kinetics assay of drP18 C50S at varying protein concentrations (10–40 μM), oxidized with different oxidant concentrations (10–400 μM) while maintaining a 1:10 protein-to-oxidant ratio, whereas a 1:1 protein-to-HOSCN ratio was used for HOSCN. (C) Thioflavin-T fluorescence kinetics assay of drP18 C50S (20 μM) oxidized with diamide (200 μM, left), $HCO_4^-$ (200 μM, center), and HOSCN (20 μM, right). Addition of 2 mM DTT at 1430 min (indicated by arrow) reduces fluorescence to baseline only in the $HCO_4^-$-treated sample, whereas the other two conditions show a gradual decrease without returning to baseline.

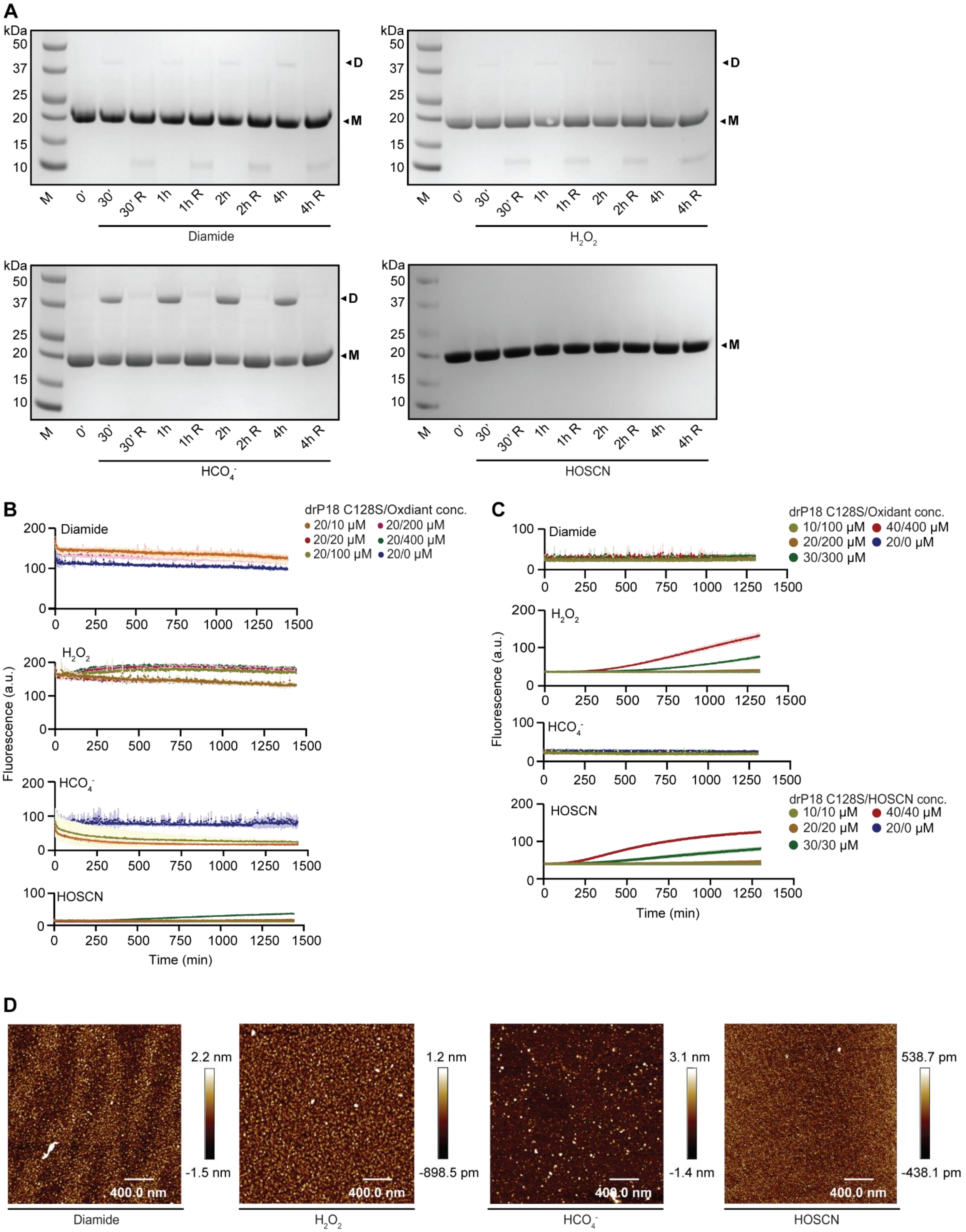
A
kDa
50
37
25
20
15
10
D
M
M 0' 30' 30' R 1h 1h R 2h 2h R 4h 4h R
Diamide
H2O2
HCO4-
HOSCN
B
C
D
drP18 C128S/Oxidant conc.
20/10 μM 20/200 μM
20/20 μM 20/400 μM
20/100 μM 20/0 μM
10/100 μM 40/400 μM
20/200 μM 20/0 μM
30/300 μM
drP18 C128S/HOSCN conc.
10/10 μM 40/40 μM
20/20 μM 20/0 μM
30/30 μM
Fluorescence (a.u.)
Time (min)
2.2 nm
-1.5 nm
1.2 nm
-898.5 pm
3.1 nm
-1.4 nm
538.7 pm
-438.1 pm
400.0 nm

**Figure S7. Oxidation of drP18 C128S with different oxidants.** (A) SDS-PAGE analysis of 20 µM drP18 C128S (M) following oxidation with 200 µM Diamide, $H_2O_2$, $HCO_4^-$ and 20 µM HOSCN over a time course of 4 h. D, dimeric species, was detected under non-reducing conditions, which was largely reduced in the presence of DTT. (B) Thioflavin-T fluorescence kinetics assay of drP18 C128S (20 µM) oxidized with different oxidants at concentrations ranging from 10–400 µM. (C) Thioflavin-T fluorescence kinetics assay of drP18 C128S at varying protein concentrations (10–40 µM), oxidized with different oxidant concentrations (10–400 µM) while maintaining a 1:10 protein-to-oxidant ratio, whereas a 1:1 protein-to-HOSCN ratio was used for HOSCN. (D) AFM characterization of drP18 C128S. Representative images of oxidized samples (24 h at 25 °C), including diamide (left), $HCO_4^-$ (center left), $H_2O_2$ (center right), and HOSCN (right). Scale bar: 400 nm.

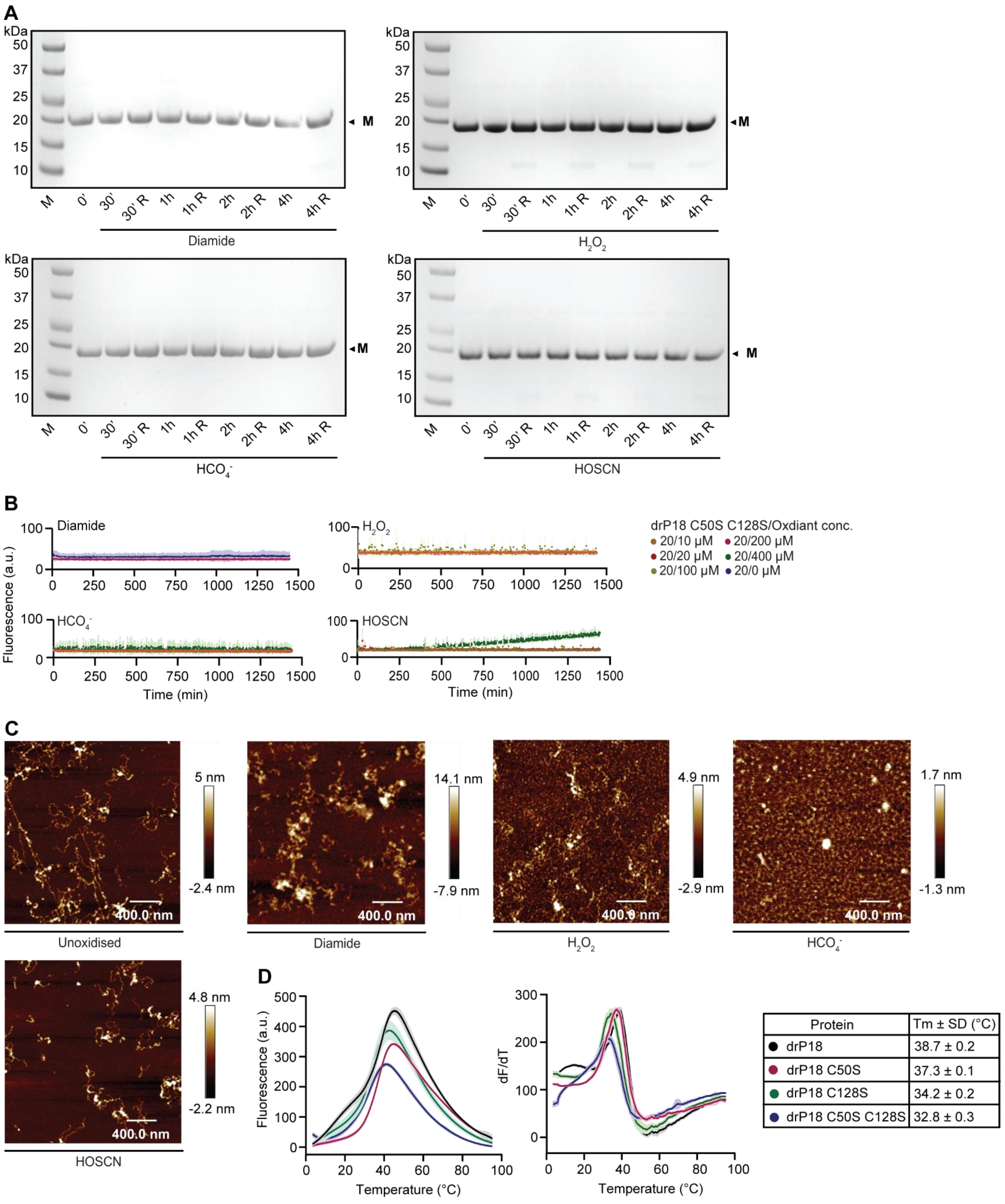


**Figure S8. Oxidation of drP18 C50S C128S with different oxidants.** (A) SDS-PAGE analysis of 20 µM drP18 C50S C128S following oxidation with diamide (200 µM), $H_2O_2$ (200 µM), $HCO_4^-$ (200 µM), or HOSCN (20 µM) over a 4 h time course. The protein remained predominantly monomeric under all conditions. (B) Thioflavin-T fluorescence kinetics assay of drP18 C50S C128S (20 µM) oxidized with different oxidants at concentrations ranging from 10–400 µM. (C) AFM characterization of drP18 C50S C128S. Samples were incubated for 24 h at 25 °C, with representative images of unoxidized (left), diamide-oxidized (center left), $HCO_4^-$ -oxidized (center right), $H_2O_2$ -oxidized (right), and HOSCN-oxidized (bottom) protein. Fibrillar structures were observed even in unoxidized samples, indicating a loss of structural integrity upon cysteine substitution. Scale bar: 400 nm. (D) Differential scanning fluorimetry (DSF) melt curves of drP18 and variants. Raw fluorescence as a function of temperature (left), first derivative (dF/dT) (right) curves used to determine melting temperatures ($T_m$). Black, drP18; pink, drP18 C50S; green, drP18 C128S; blue, drP18 C50S C128S. Table showing the melting temperature ($T_m$) of different protein variants.

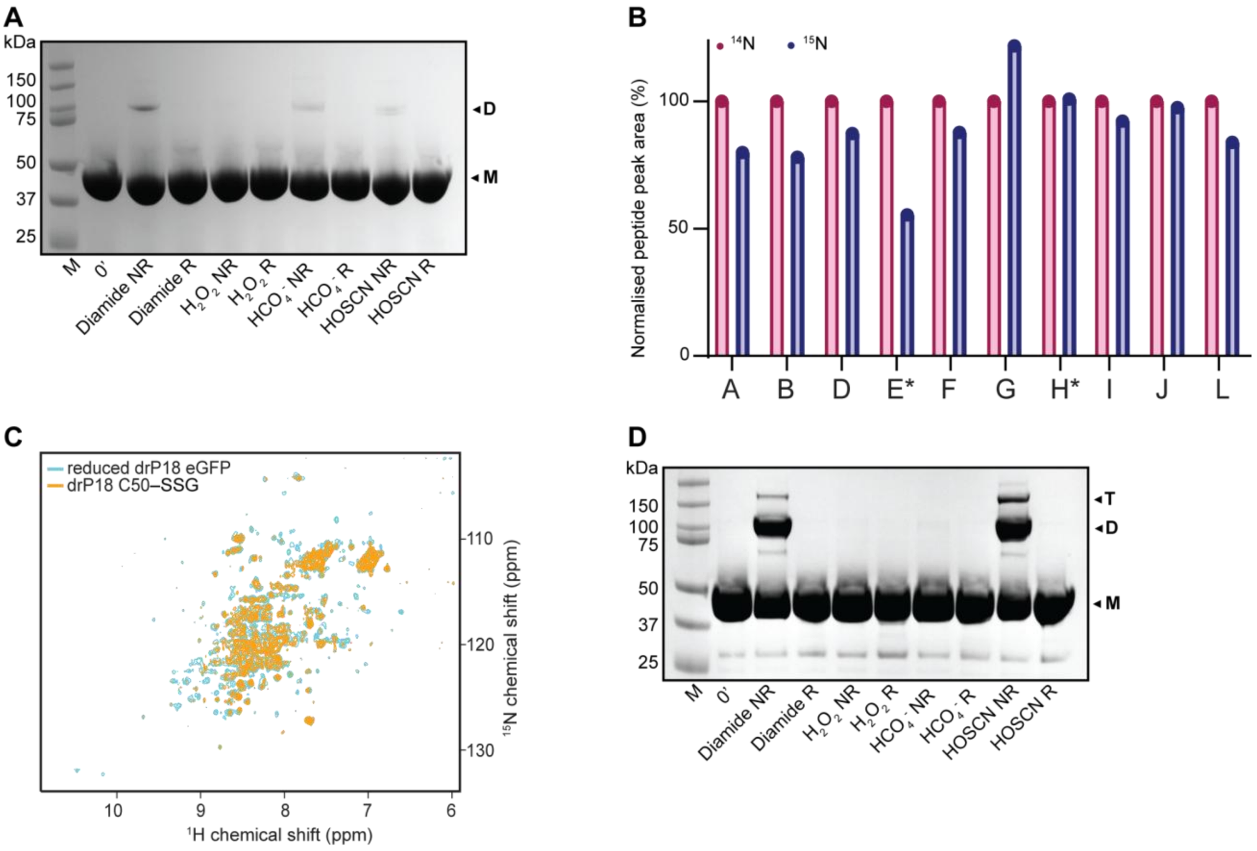


**Figure S9. Oxidation and glutathionylation of drP18 eGFP.** (A) SDS-PAGE analysis of recombinantly expressed 20 μM drP18 eGFP (M) shows the formation of D, dimers following oxidation with 200 μM diamide, $H_2O_2$, and $HCO_4^-$, and 20 μM HOSCN over a 4 h time course at 25 °C. DTT addition reverses the disulfide-linked bands. (B) Glutathione-treated $^{15}N$-labelled drP18 eGFP was mixed with reduced $^{14}N$-labelled drP18 eGFP as an internal reference standard, alkylated with NEM, digested with trypsin, and analyzed by LC-MS. Peptide peak areas were quantified and normalized to the corresponding $^{14}N$ reference peptide. Peak area intensity reflects the relative abundance of the corresponding peptide species. Cysteine-containing peptides are indicated by an asterisk (*) (full tryptic digest is shown in Fig. S4; the corresponding peptide sequence and theoretical m/z values are shown in Table S2). The C50-containing peptide (E) exhibited a pronounced reduction in NEM-labelled $^{15}N$ signal relative to the corresponding $^{14}N$ reference peptide, indicating cysteine occupancy by glutathione and confirming predominant *S*-glutathionylation at C50. (C) $^{1}H^{15}N$ HSQC solution NMR spectrum of drP18 eGFP in the reduced state (cyan) and after *S*-glutathionylation (orange). Additional resonances from the eGFP tag are present, but chemical shifts show similar changes compared to the untagged version upon glutathionylation. (D) SDS-PAGE analysis of 20 μM drP18 eGFP C50–SSG (M) following oxidation with 200 μM diamide, $H_2O_2$, or $HCO_4^-$, or 20 μM HOSCN, over a 4-h time course at 25°C. T, Trimers and D, dimers, were observed following diamide and HOSCN treatment, whereas faint dimer bands were detected following $H_2O_2$ and $HCO_4^-$ treatment. Addition of DTT reversed the disulfide-linked oligomeric species.

| **Peptide (isotope mix)** | **Parent ion (charge state)** | **Fragment ion** |
|---|---|---|
| LGCPSIAETLLQAGADPNVR-LGCPSIAETLLQAGADPNVR disulfide ($^{14}N$-$^{14}N$) | 1013.15 m/z (+4) | $y_4$: 485.28 m/z |
| LGCPSIAETLLQAGADPNVR-LGCPSIAETLLQAGADPNVR disulfide ($^{14}N$-$^{15}N$) | 1019.40 m/z (+4) | $y_4$: 485.28/493.28 m/z |
| LGCPSIAETLLQAGADPNVR-LGCPSIAETLLQAGADPNVR disulfide ($^{15}N$-$^{15}N$) | 1025.65 m/z (+4) | $y_4$: 493.28 m/z |
| EGHLDVVQFLVTHCVTQPFLANAK-EGHLDVVQFLVTHCVTQPFLANAK disulfide ($^{14}N$-$^{14}N$) | 1067.42 m/z (+5) | $y_7$: 760.44 m/z |
| EGHLDVVQFLVTHCVTQPFLANAK-EGHLDVVQFLVTHCVTQPFLANAK disulfide ($^{14}N$-$^{15}N$) | 1073.82 m/z (+5) | $y_7$: 760.44/ 769.44 m/z |
| EGHLDVVQFLVTHCVTQPFLANAK-EGHLDVVQFLVTHCVTQPFLANAK disulfide ($^{15}N$-$^{15}N$) | 1080.22 m/z (+5) | $y_7$: 769.44 m/z |
| LGCPSIAETLLQAGADPNVR-EGHLDVVQFLVTHCVTQPFLANAK disulfide ($^{14}N$-$^{14}N$) | 939.07 m/z (+5) | $y_7$: 728.37 m/z * |
| LGCPSIAETLLQAGADPNVR-EGHLDVVQFLVTHCVTQPFLANAK disulfide ($^{15}N$-$^{14}N$) | 944.07 m/z (+5) | $y_7$: 739.37 m/z * |
| LGCPSIAETLLQAGADPNVR-EGHLDVVQFLVTHCVTQPFLANAK disulfide ($^{14}N$-$^{15}N$) | 945.47 m/z (+5) | $y_7$: 728.37 m/z * |
| LGCPSIAETLLQAGADPNVR-EGHLDVVQFLVTHCVTQPFLANAK disulfide ($^{15}N$-$^{15}N$) | 950.47 m/z (+5) | $y_7$: 739.37 m/z * |

**Table S1.** The m/z values for the parent ions and the daughter fragment ions that were used to quantify each peptide in LC-MS/MS experiments. *$y_7$ fragment ion is from LGCPSIAETLLQAGADPNVR peptide.

| Peptide | $^{14}$N peptide m/z values | | | | $^{15}$N peptide m/z values | | | |
|---|---|---|---|---|---|---|---|---|
| | +1 | +2 | +3 | +4 | +1 | +2 | +3 | +4 |
| GAMAE DTALDR | 1149.52 | 575.26 | | | 1163.52 | 582.26 | | |
| LSTAAAI GDLME VEQTLQ SNVNV NEK | *2774.39* | 1387.70 | | | *2806.39* | 1403.70 | | |
| TALQVM K | 790.45 | 395.73 | | | 799.45 | 400.23 | | |
| LGC(NE M)PSIAE TLLQAG ADPNV R | *2150.09* | 1075.55 | | | *2175.09* | 1088.05 | | |
| DPILGL TVIHDA AR | 1490.84 | 745.92 | 497.62 | | 1509.84 | 755.42 | 503.95 | |
| DGYLDT LHVLAQ NGADV NLLDND GNLPLH LAAR | *3527.79* | 1764.40 | 1176.60 | 882.70 | *3572.79* | 1786.90 | 1191.60 | 893.95 |
| EGHLDV VQFLVT HC(NEM )VTQPF LANAK | *2791.43* | 1396.22 | 931.15 | 698.61 | *2823.43* | 1412.22 | 941.81 | 706.61 |
| GYTPR | 593.31 | 297.16 | | | 601.31 | 301.16 | | |
| DLAFMH QK | 989.49 | 495.25 | 330.50 | | 1001.49 | 501.25 | 334.50 | |
| TVEWLE SIASLQ SSQR | 1833.94 | 917.47 | | | 1855.94 | 928.47 | | |

**Table S2.** Theoretical m/z values of $^{14}$N- and $^{15}$N-labelled peptides used for peptide mass spectrometry analysis. Expected m/z values are shown for the +1 to +4 charge states of each peptide. Cysteine residues modified with *N*-ethylmaleimide (NEM) are indicated as …C(NEM). The table includes the cysteine-containing peptides used to identify the site of glutathionylation shown in the corresponding mass spectrometry figure. The charge states used for peptide quantification are shown in normal font, whereas charge states with m/z values outside of the range of the mass spectrometer (>2000 m/z) are shown in italics.
